# An integrative and multi-indicator approach for wildlife health applied to an endangered caribou herd

**DOI:** 10.1101/2023.02.01.526641

**Authors:** Xavier Fernandez Aguilar, Lisa-Marie Leclerc, Kugluktuk Angoniatit Association, Ekaluktutiak Hunters & Trappers Organization, Olokhaktomiut Hunters & Trappers Committee, Fabien Mavrot, Amelie Roberto-Charron, Matilde Tomaselli, Gabriela Mastromonaco, Anne Gunn, Mathieu Pruvot, Jamie L. Rothenburger, Niroshan Thanthrige-Don, Elham Zeini Jahromi, Susan Kutz

## Abstract

Assessing wildlife health in remote regions requires a multi-faceted approach that may include convenience samplings and the involvement of local communities. Combining data from hunted and captured caribou, we assessed the value of multiple indicators for understanding the health of the endangered Dolphin and Union caribou herd in Arctic Canada. We documented health determinants (infectious diseases and trace elements), processes (cortisol, pathology), and health outcomes (pregnancy and body condition). During a recent period of steep population decline our results suggested relatively good body condition and pregnancy rates and decreasing levels of stress, along with low adult cow survival. We identified multiple determinants of health as potential contributors to reduced survival, including *Brucella* suis biovar 4, *Erysipelothrix rhusiopathiae* and lower hair trace minerals. This integrative approach that drew on multiple data sources has provided unprecedented knowledge on the health in this herd and highlights the value of documenting individual animal health to understand causes of wildlife declines.

## Introduction

Highly seasonal and high latitude regions such as the Arctic are especially vulnerable to the effects of accelerating climate warming, such as changes in plant composition and phenology (Myers-Smith et al. 2011; Post et al. 2018), shifting wildlife and parasite distributions and communities (Post et al. 2009; Kafle et al. 2020), and altered host-parasite interactions (Kutz et al. 2005). These changes can exceed the resilience of wild animal populations, resulting in increased morbidity and mortality (Hunter et al. 2010; Kutz et al. 2015; Tomaselli et al. 2018). The remoteness of the Arctic makes health surveillance and investigations especially challenging. Multi-faceted approaches are required to track wildlife health, identify emerging threats, and implement wildlife conservation measures (Kutz and Tomaselli 2019).

The majority of migratory tundra caribou (*Rangifer tarandus*) herds in North America have declined since the early 2000s, with only a few herds currently stable or increasing (Russell et al. 2018). About 2.6 to 4.7 million caribou have disappeared from the Arctic in the last two decades, with an average herd population decline of 71% ± 5.9SE from their peak (11 herds with sufficient data), and some of the herds have plummeted by more than 90% (Gunn and Russell 2017; Russell et al. 2018). Although both traditional and scientific knowledge documented historical fluctuations of tundra caribou (Zalatan et al. 2006; Hanke et al. 2022), several herds are now below any historically documented population levels and show few signs of recovery (Russell et al. 2018).

The magnitude of the migratory tundra declines is in stark contrast with our poor understanding of the drivers of the declines. While climate change may be an overarching driver of caribou population trends (Mallory and Boyce 2018), the specific mechanisms causing the declines are poorly understood. This is especially true for the role of nutrition, disease and parasites, and how these health determinants interact with each other (Macbeth and Kutz 2019). Demographic indicators of caribou populations are regularly monitored, yet these alone are insufficient to understand the underlying drivers of declines. Deeper and comprehensive health investigations are necessary to improve our understanding on how climate change is altering the Arctic systems, identify health and demographic drivers, anticipate threats and, if possible, mitigate them through evidence-based management decisions.

Recent partnerships with Indigenous communities have progressed to establish health surveillance systems through sustained and standardized community-based wildlife health monitoring programs. These programs greatly expand on the sporadic sampling derived from caribou captures and collaring operations (Peacock et al. 2020; Tomaselli 2022). In fact, community-based surveillance and local knowledge can detect shifts in health and drivers that may not be detected by other classical monitoring methods that are more restricted by budget or the frequency and seasonality of observations (Tomaselli et al. 2018; Kutz and Tomaselli 2019). Peacock et al. proposed an integrative framework that incorporates different health metrics informed by local and scientific knowledge to quantitatively assess changes in Arctic wildlife health. This framework is based on benchmarks that are yet to be defined for caribou and baseline data are much needed (Peacock et al. 2020).

The complexity of multiple factors influencing health is conceptually represented by the Determinants of Health model, which was conceived for human health and has been adapted to wildlife (Wittrock et la., 2019). These conceptual models acknowledge the accumulative effects of a wide range of determinants, yet do not clearly differentiate between causes and consequences. They also do not include previously discussed concepts on wildlife health such as the differentiation between determinants and outcomes (Ryser-Degiorgis 2013). With the aim to operationalize health assessments in wildlife, a novel framework was further elaborated upon in Pruvot et al., by classifying health indices according to the type of information they provide (Pruvot et al.).

The goal of our research was to build on existing wildlife health frameworks to evaluate the significance of various individual animal health indices for understanding the health of an endangered tundra caribou herd, the Dolphin and Union herd (*Rangifer tarandus groenlandicus* x *R. t. pearyi*). This caribou herd has experienced recent steep population declines and since 2015, monitoring efforts including capture and collaring events and community-based surveillance have been intensified, generating an unprecedented amount of health information on this herd. We compiled this information and applied a structured approach to classify and analyze different health indices into health determinants, health processes, and health outcomes. With this study, we aim to share experiences on this novel approach, identify mechanisms and drivers of the recent caribou herd decline, identify relevant health indicators and further surveillance needs on caribou health, and critically assess the combination of community-based sampling and sampling from captured animals for health investigations.

## Material and Methods

### Study area, population, and sampling methods

The Dolphin and Union (DU) caribou is a migratory tundra caribou herd endemic to the Canadian Arctic (Nunavut and Northwest Territories), and is considered a separate conservation and management unit because its unique genetics, morphology, and ecology (COSEWIC 2017). This herd has been steadily declining since approximately the 1990s-2000s, with the last estimate of 3,815 in 2020 (95% CI: 2,930–4,966) (Campbell et al. 2021); it is currently considered endangered in Canada (COSEWIC). The biggest herd decline (78%) was documented between 2015 and 2018 (Leclerc and Boulanger 2020). Detailed information is provided in Supplementary Information S1.

There are three Inuit communities, Kugluktuk, Cambridge Bay and Ulukhaktok, that harvest DU caribou at different locations along its seasonal migration. A collaborative health monitoring program for this herd was initiated in 2015 and has been facilitated through the Kutz Research Group, the Hunter and Trappers associations (HTO/HTCs) from Kugluktuk, Cambridge Bay, Ulukhaktok and the Governments of Nunavut and Northwest Territories. It includes two different sources of convenience sampling, caribou hunted for subsistence or recreational purposes (guided hunting) and sampling from captured animals for monitoring purposes by the Government of Nunavut. The involvement of Inuit communities in sample and data collection was coordinated through HTO/HTCs and started at different times depending on funding sources (Table S1).

Captures were performed by helicopter to deploy satellite collars on adult females (Leclerc and Boulanger 2018). In both community-based sampling and live-captured caribou, sampling methods were standardized following modified protocols of the CircumArctic Rangifer Monitoring and Assessment Network (CARMA) (Kutz et al. 2013). The specific sample list collected, methods of management and storage of samples and research permits are specified in Supplementary Information S1.

Health data from the DU herd from previous periods were available through the CARMA network (Don Russell). This included data of caribou body condition and health from government surveys between 1987 and 1991 (Gunn et al. 1991), and data from a research study that took place between 2001 to 2003 on caribou harvested by local hunters in a similar way as hunters harvest animals for subsistence (Hughes et al. 2009).

### Sample Analyses and health indices

*We* applied a health framework that classifies health indicators into health determinants, health processes and health outcomes (Figure 1, Table 1). This framework elaborates upon previous concepts on health assessments and surveillance (Ryser-Degiorgis 2013; Wittrock et al. 2019), and the theory behind this framework is described in depth in (Pruvot et al.). Briefly, health determinants can be considered as extrinsic or intrinsic factors that can promote or affect health (Macbeth and Kutz 2019; Wittrock et al. 2019). Health processes are the animal’s physiological, pathological or behavioural processes that occur in response to determinants, termed ‘host responses’ here. These responses, e.g. stress hormones or acute phase proteins, are often not specific to a single determinant and therefore offer broader information about possible drivers of health. Finally, health outcomes are the result of the interaction of determinants of health and the resilience of caribou, and include those parameters that are associated with fitness, such as reproduction or survival. For capital breeders, such as caribou, energy stores are critical and can be considered as a proxy for fecundity (Parker et al. 2009; Albon et al. 2017), thus in this work we used this indicator as a health outcome.

**Figure 1.**
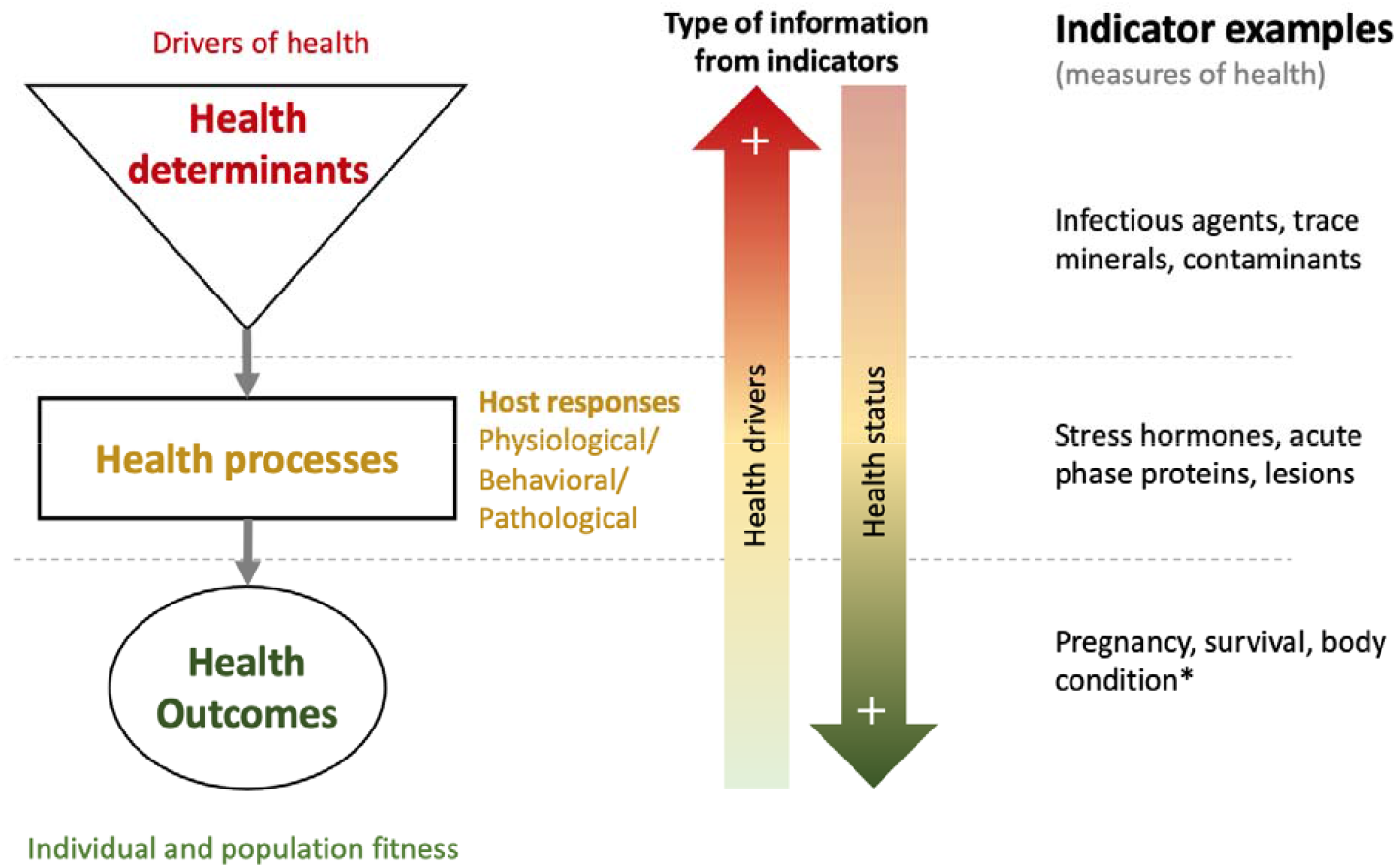
Classification and examples of health indicators as health determinants, health processes (host responses to determinants) and health outcomes showing a directional relationship in a gradient of information between health drivers and health status. This diagram depicts the main relationships between indicator class types but does not intend to encompass the whole theoretical complexity behind this framework, *body condition may be considered as a health process as well, see full text for justifications.

**Table 1.**
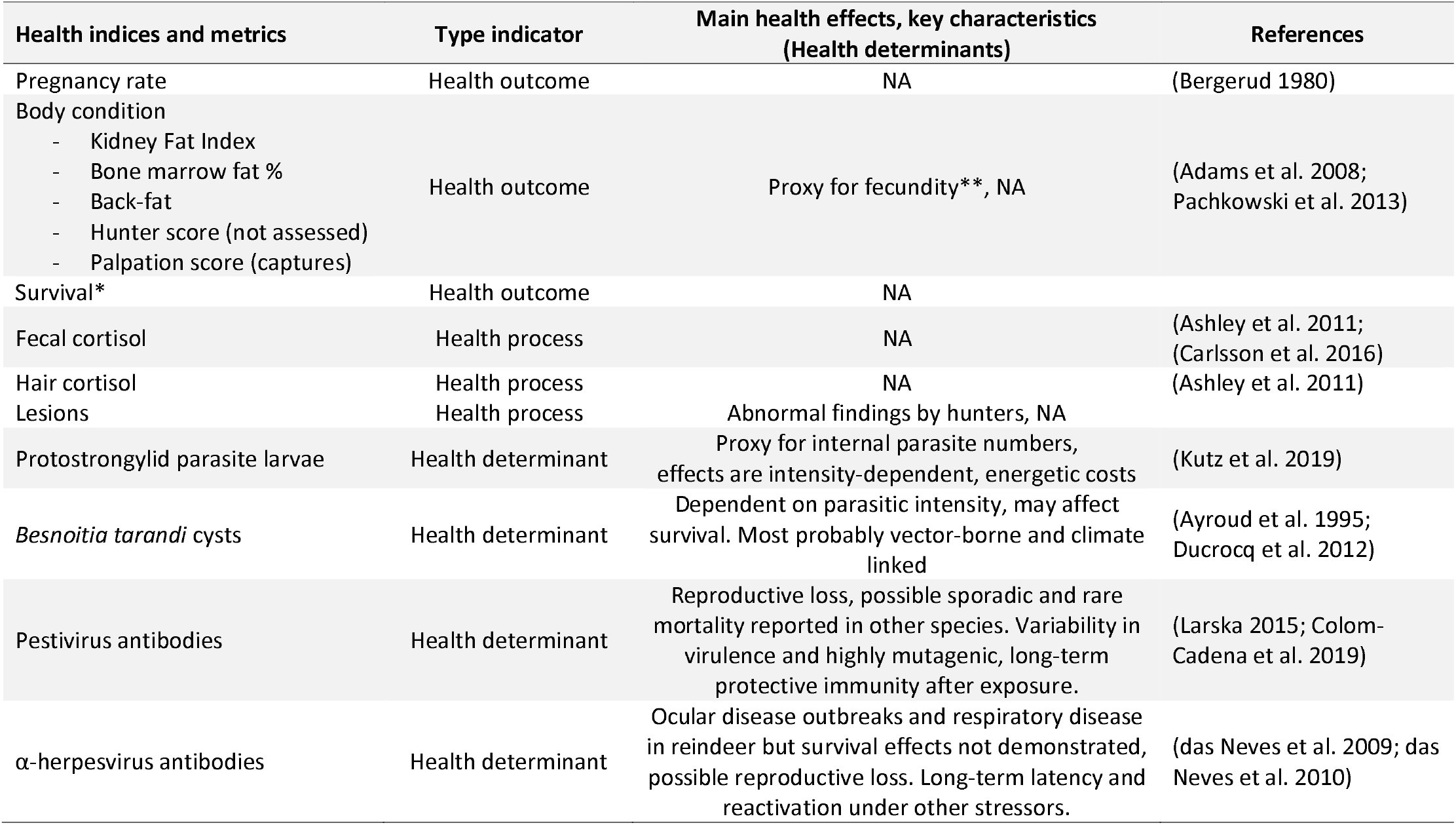
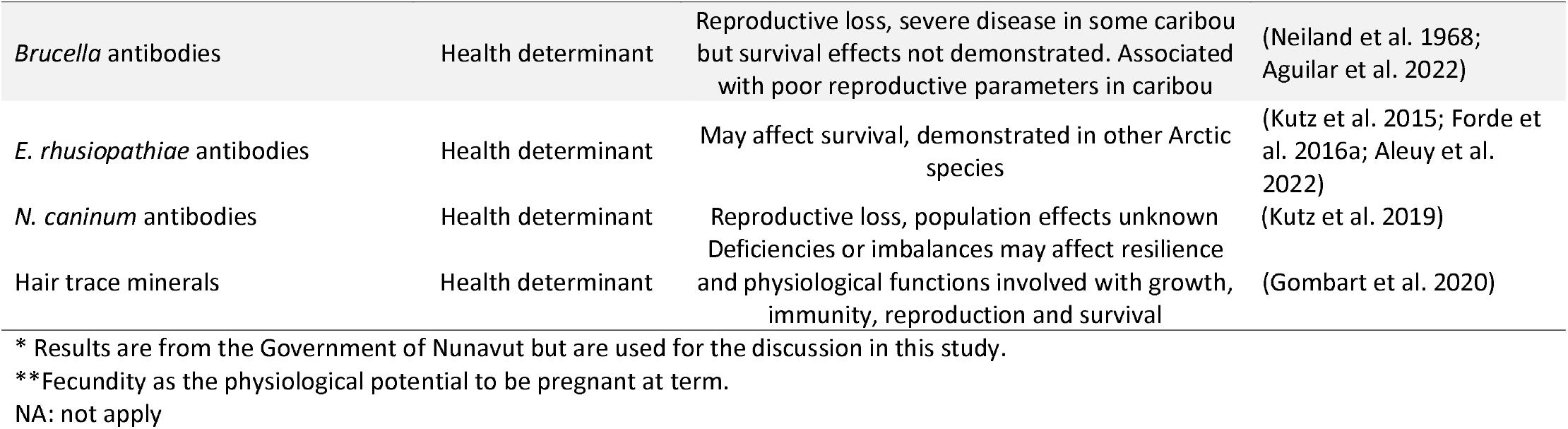
Indices used in this study for the health assessment of the Dolphin and Union caribou herd. The effects on health of determinants are based on available knowledge on caribou or related species, or the known pathobiology of the infectious agents.

| Health indices and metrics | Type indicator | Main health effects, key characteristics<br>(Health determinants) | References |
| --- | --- | --- | --- |
| Pregnancy rate | Health outcome | NA | (Bergerud 1980) |
| Body condition | Health outcome | Proxy for fecundity**, NA | (Adams et al. 2008;<br>Pachkowski et al. 2013) |
| - Kidney Fat Index |  |  |  |
| - Bone marrow fat % |  |  |  |
| - Back-fat |  |  |  |
| - Hunter score (not assessed) |  |  |  |
| - Palpation score (captures) |  |  |  |
| Survival* | Health outcome | NA |  |
| Fecal cortisol | Health process | NA | (Ashley et al. 2011;<br>(Carlsson et al. 2016) |
| Hair cortisol | Health process | NA | (Ashley et al. 2011) |
| Lesions | Health process | Abnormal findings by hunters, NA |  |
| Protostrongylid parasite larvae | Health determinant | Proxy for internal parasite numbers, effects are intensity-dependent, energetic costs | (Kutz et al. 2019) |
| *Besnoitia tarandi* cysts | Health determinant | Dependent on parasitic intensity, may affect survival. Most probably vector-borne and climate linked | (Ayroud et al. 1995;<br>Ducrocq et al. 2012) |
| Pestivirus antibodies | Health determinant | Reproductive loss, possible sporadic and rare mortality reported in other species. Variability in virulence and highly mutagenic, long-term protective immunity after exposure. | (Larska 2015; Colom-Cadena et al. 2019) |
| α-herpesvirus antibodies | Health determinant | Ocular disease outbreaks and respiratory disease in reindeer but survival effects not demonstrated, possible reproductive loss. Long-term latency and reactivation under other stressors. | (das Neves et al. 2009; das Neves et al. 2010) |
| *Brucella* antibodies | Health determinant | Reproductive loss, severe disease in some caribou but survival effects not demonstrated. Associated with poor reproductive parameters in caribou | (Neiland et al. 1968; Aguilar et al. 2022) |
| *E. rhusiopathiae* antibodies | Health determinant | May affect survival, demonstrated in other Arctic species | (Kutz et al. 2015; Forde et al. 2016a; Aleuy et al. 2022) |
| *N. caninum* antibodies | Health determinant | Reproductive loss, population effects unknown | (Kutz et al. 2019) |
| Hair trace minerals | Health determinant | Deficiencies or imbalances may affect resilience and physiological functions involved with growth, immunity, reproduction and survival | (Gombart et al. 2020) |

\* Results are from the Government of Nunavut but are used for the discussion in this study.
\*\*Fecundity as the physiological potential to be pregnant at term.
NA: not apply

Health outcomes provide perhaps the most relevant information needed for managers to anticipate the herd trajectory. Health outcomes alone, however, do not inform about the underlying drivers and processes leading to the health outcome. Parameters that measure health processes may provide some insights into the health status and may (or may not) inform on the determinants that triggered that response. Determinants of health, on the other hand, are those processes that are driving the health outcome and could be a possible target of management. In this study we first focus on the health outcomes, and then use the information provided by metrics on responses and potential determinants to identify what drivers may be operating on the herd’s health.

### Health outcomes

Health outcomes included body condition, pregnancy status and data on survival (Table 1). Body condition metrics in harvested animals included the percentage of metatarsus marrow fat, Kidney Fat index (KFI), back-fat depth and a hunter qualitative assessment following the CARMA protocol (Adams et al. 2008). In live-captured animals, a body condition score that range from 1 to 12 was derived from the palpation of different body parts (Supplementary Information S2).

Pregnancy status of caribou sampled in early spring (April and May) was inferred from fecal progesterone levels, following previously described protocols, and establishing within-population thresholds for non-pregnant females at <200ng/g and pregnant females at >700 ng/g of wet feces (Morden et al. 2011).

Where data collection methods were comparable, we compared our results on KFI and pregnancy rates in adult females in early spring with data from the DU caribou herd in 1987-1991 and 2001-2003.

### Health processes (host responses to determinants)

Physiological stress was measured indirectly using cortisol and its metabolites in hair and feces of the animals. Fecal stress hormone levels reflect physiological status from approximately the last 24-48 hours before sample collection. Hair values largely reflect the accumulation of cortisol during the hair growth period from late spring to fall, representing conditions over several months during the previous summer and fall, and possibly also reflecting stressors during the non-hair growing season (Rakic et al.). Cortisol concentrations (hair from the neck) and corticosteroid metabolites (feces) were quantified by enzyme immunoassay (Carlsson et al. 2016; Supplementary Information S3). Hair cortisol from 2015-2017 and 2018-2021 was analyzed in different laboratories (Supplementary Information S3).

### Health Determinants

For the assessment of pathogen exposure, blood collected with filter paper from both captured and hunted animals was eluted for antibody detection (Curry et al. 2011). The presence of antibodies against selected pathogens, including α-herpesvirus, pestivirus, *Brucella, Erysipelothri× rhusiopathiae* and *Neospora caninum*, was tested assuming the 1:10 dilution of filter paper eluates (Curry et al. 2011) (see Table S2 for test and laboratory information). The selection of pathogen assays was based on previous knowledge from this herd or other tundra caribou herds (Carlsson et al. 2019; Kutz et al. 2019).

To estimate the intensity of infection of *Besnoitia tarandi, we* used the maximal density of bradyzoite-containing cysts (mdc) from the anterior aspect of the mid-third portion of the metatarsal skin region (Ducrocq et al. 2012)(Supplementary Information S4). We used the modified Baermann technique for the quantification of protostrongylid larvae from frozen feces (Kafle et al. 2017), but larvae of *Parelaphostrongylus andersoni* and/or *Varestrongylus eleguneniensis* were not differentiated (Schindelin et al. 2012).

We determined the element concentrations in the hair from the neck collected from 2018 to 2021 using inductively coupled plasma mass spectrometry (ICP-MS) at the Alberta Centre for Toxicology. The methods used and the panel of trace minerals and contaminants analyzed are described in detail in Supplementary Information S5.

### Analyses for ancillary data

Caribou sampled were confirmed to belong to the Dolphin and Union herd by genetic analyses or, in the absence of these data, on sampling location and morphological traits described by harvesters (Leclerc and Boulanger 2018)(Supplementary Information S6). The age of caribou was assessed by analyzing the cementum annuli from an incisor or by the tooth eruption pattern of the incisors (Adams et al. 2008). Age class was assigned as calf for <12 months, yearling for 13-24 months, subadult for 25-36 months and adult for 37 months or greater.

### Data analyses and assessments

*We* explored the distribution of data and its structure in relation to possible sources of covariation. Most samples were from adult caribou (88.2% of the samples, n=247) and, therefore, we only used this age category for our assessments. Because the dataset was highly unbalanced between years regarding the method and season of sampling or sex, and continuous variables were not normally distributed, we used univariate tests and subsets of data for statistical analyses. Data derived from hair samples were corrected to the corresponding hair-growing year, thus samples collected from January through to spring were re-assigned to the previous calendar year when the hair would have been growing.

We used a subset of data with adult DU caribou sampled in the same year and the same season (April and May 2018, n=86) to evaluate differences between the two sampling sources, hunted caribou and captured animals. For this purpose, we used Fisher’s Exact Tests and Chi-square tests for independence to assess differences by sampling source in pregnancy status and exposure to pathogens; Wilcoxon-Mann-Whitney tests to assess differences in fecal/hair cortisol, and hair trace minerals. Body condition was recorded differently for live (palpation) versus harvested (KFI, marrow fat, back-fat) animals and therefore differences of body condition by sampling source were not assessed.

For hair cortisol analyses, we excluded results corresponding to 2015 (hair samples collected in the fall 2015 and spring 2016) because were analyzed with a different test and laboratory. Differences between years were assessed in the overall sample set of adult caribou in early spring (April and May), selecting females or a specific sampling source depending on the number of observations and whenever these groups were not significant in previous assessments. We used Fisher’s Exact Tests and Chi-square tests for independence when variables were categorical and non-parametric Kruskal–Wallis and Wilcoxon-Mann-Whitney tests when variables were continuous.

Pregnancy rates and body condition of adult females in spring were compared among sampling periods (2015-2019, 2001-2003 and 1987-1992) using Chi-square and Kruskal-Wallis tests, respectively.

All analyses and data figures were done using the R software version 4.1.0 (R Development Core Team 2017), and significance level were set at 0.05 in all tests.

## Results

### Dolphin and Union caribou sampling

A total of 298 caribou were sampled between 2015 and 2021 (Table S1 and Figure 2). Eighteen animals were eliminated from the dataset after genetic analyses revealed they had barren-ground genetics (>50% genetic profile, Supplementary Information S6). Sample submissions varied by year and in part due to availability of specific funding (subsistence hunts from 2015 to 2017) and disruption of sampling during the COVID-19 pandemic (2020). Sport hunts on the DU caribou ceased in 2019 in response to low population estimates, and samples from captured caribou (adult females) were only available for the years when the Government of Nunavut were deploying GPS collars.

**Figure 2.**
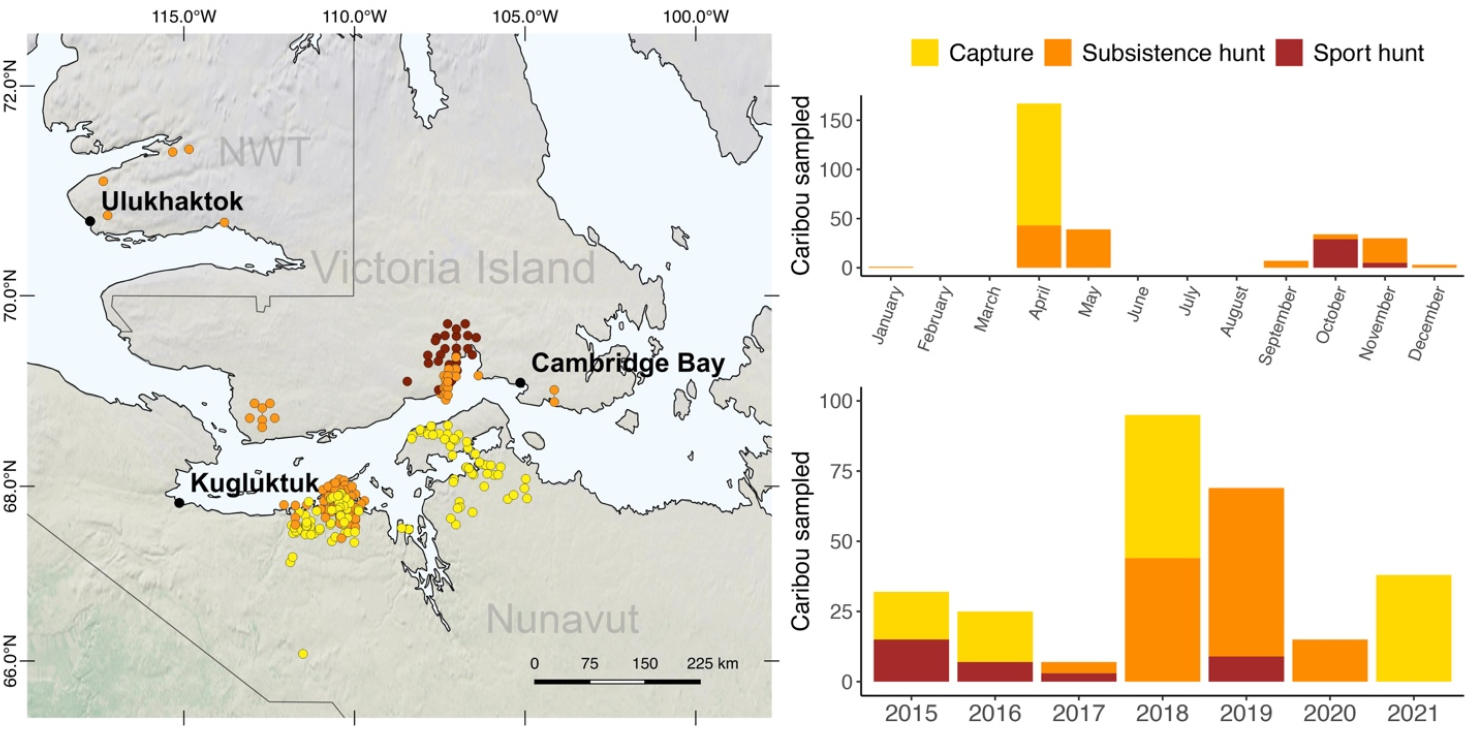
Left: locations of the Dolphin and Union (DU) caribou sampled since 2015 in Nunavut and Northwest Territories (Canada), and the three communities that are involved in harvester-based sampling. Right: Number of DU caribou sampled over the years by sampling source. The map was produced using QGIS (QGIS Development Team 2016, Las Palmas 2.18.15; QGIS Geographic Information System. Open-Source Geospatial Foundation Project).

Most of the samples were collected in early spring (73% from mid-April to mid-May), and less commonly, in the fall (25% from mid-September to mid-November). These two periods are when the DU herd is closer to communities and when most of the harvests and captures (spring only) happen. This seasonal sampling helped to standardize data within and across years, but limits insight into the calving and summer period.

### Results on health outcomes

The average pregnancy rate in adult caribou sampled in early spring for the period 2015-2021 was 86.8% (95% CI 80.9-91.1). In 2018, pregnancy rates were significantly lower in hunted 67.9% (95% CI 49.3-82.1) vs captured 93.7% (95% CI 83.2-97.9) caribou (n= 76, p=0.007 Fisher’s exact test). For the period 2015-2019, the average pregnancy rate of hunted adult females in early spring (78.7%; 95% CI: 61.8-83.1) was similar to that documented for the period 1987-1991 (76.2%; 95% CI: 67.3-82-7), but significantly higher than for the period 2001-2003 (57%; 95% CI: 46.5-67.5, *X*^2^ (df= 2, n= 209) = 8.8, p=0.01) (Figure 3).

**Figure 3.**
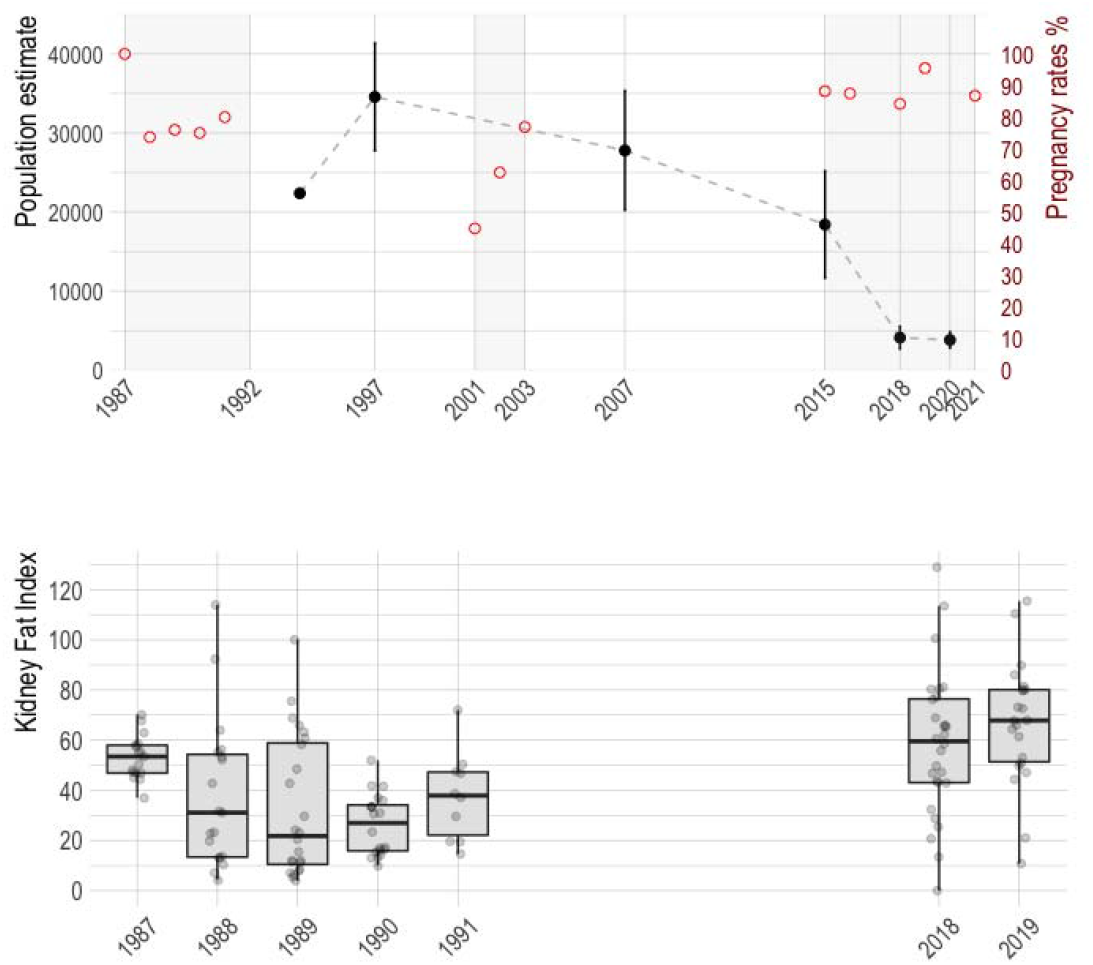
Upper chart: black dots and dashed line indicate population trends of the Dolphin and Union caribou herd (left axis), according to the Government of Nunavut population aerial surveys (Leclerc and Boulanger 2018; Leclerc and Boulanger 2020; Campbell et al. 2021). Range black lines on the dots are the standard error of the estimates. Red dots indicate the pregnancy rate estimated for that year (right axis), in three different time periods highlighted with a grey background. These pregnancy rates include estimates from adult females in early spring obtained in this study (hunted caribou: 2018 and 2019 and captured caribou: 2015, 2016, 2018, 2021), previous government surveys (hunted caribou 1987-1991) and research works (hunted-caribou 2001-2003). Lower chart: distribution of Kidney Fat Index by year, when available, from the same dataset as the upper chart.

**Figure 4.**
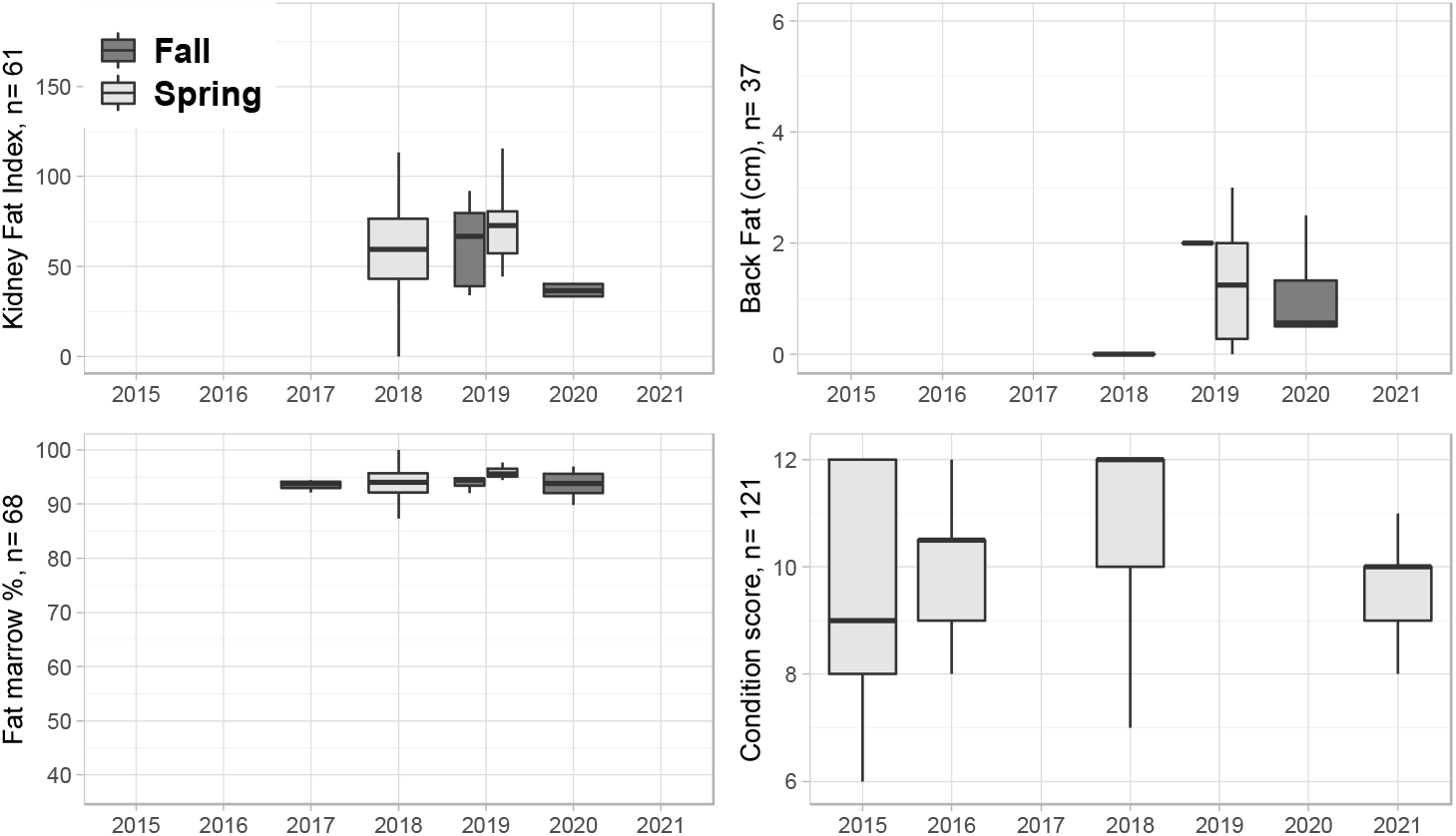
Distribution and number of observations of four different body condition metrics in adult Dolphin and Union caribou females sampled in between 2015 and 2021. Data points are overlaid on the boxplots to show sample size by year. Kidney Fat index, Back-Fat and Fat marrow % are metrics from hunted caribou and condition score from captured caribou. The hunter qualitative assessment of body condition is not represented in this figure because it had low number of observations.

The qualitative assessment of body condition by hunters and back-fat measurement were not consistently recorded on datasheets. Therefore, body condition data were mostly represented by the condition score (caribou captures) and by Kidney Fat Index and Fat marrow from females harvested at early spring (April and May) (Figure 4). Differences between years were only assessed for the condition score in captured caribou because it was the metric with the most years of data for adult females sampled during the same season. The distribution of the condition score differed significantly by years in a Kruskal Wallis H test *X*^2^ (df= 3, n= 121) = 23.2, p<0.01, with a higher median value in 2018 (12.0) than previous (2015:9.0; 2016:10.5) and later years (2021:10) (Figure 4).

Among adult females sampled in early spring, KFI median values in 2018-2019 (KFI:64.8) were statistically higher than in 1987-1991 (n= 90, KFI: 34.7), H (df= 1, n= 140) = 28.06, p<0.001 (Figure 4).

### Results on host response metrics

Data were assessed separately when hair was analyzed in different laboratories and methods (2015 vs 2017-2020). In adult caribou, the median hair cortisol value in 2015 was 14.7 pg/mg (min: 9.5-max: 20.2) and for the period 2017-2020 it was 6.7 pg/mg (min:1.0-max: 24.8). Internal research with a blinded interlaboratory comparison indicated that a 0.64 average conversion factor could be applied to compare results from the two methods/laboratories (Rakic 2022), which would still lead to a higher hair cortisol, 9.4 pg/mg in 2015, vs 8.8 pg/mg in 2017 and 4.4 pg/mg in 2020 (Figure 5).

**Figure 5.**
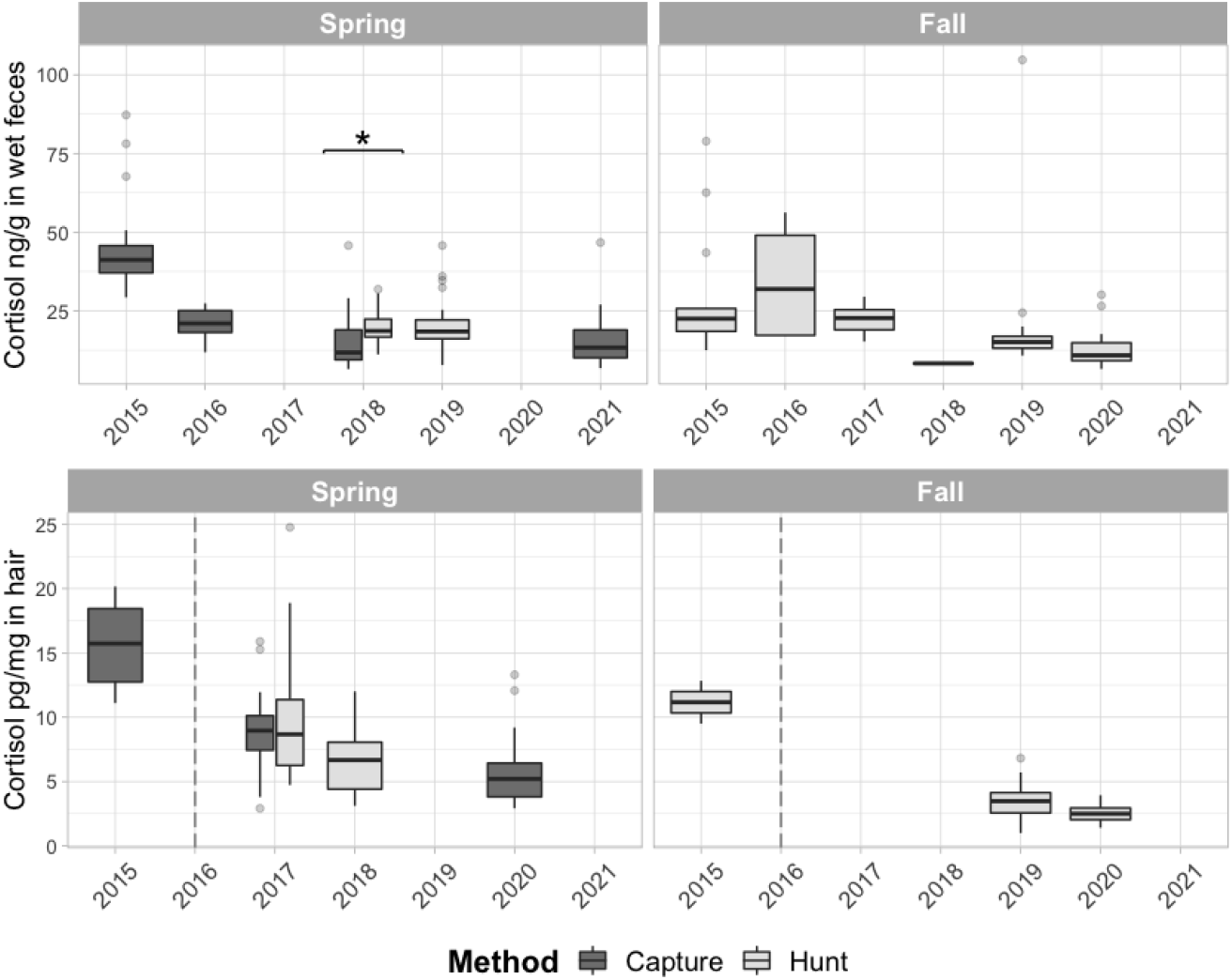
Trends of cortisol concentrations in feces from adult Dolphin and Union caribou in feces (upper panel) and in hair from the neck (lower panel), showed by season and by the sampling method used to obtain the samples. Results of hair cortisol are plotted by the year in which the hair was grown. The dashed lined in the lower panel graphs separates results that were analyzed in different laboratories and methods (2015 vs 2017-2020). Significant differences between the method of sampling are indicated with an asterisk.

We found no significant differences in hair cortisol between harvested vs captured, or sex, using a subset of data from early spring 2018. It was not possible to assess the differences of hair cortisol between sampling sources (harvest vs capture) in other years because it was confounded with season (Figure 5). Within data from 2017-2020, hair cortisol in early spring was significantly lower in later years for both captured (2017 vs 2020: n=89, Z=-5.38, p<0.01), and hunted (2017 vs 2018: n=61, Z=-3.03, p<0.01) adult caribou (Figure 5).

The overall median value for fecal cortisol in adult caribou was 17.73 ng per g of wet feces (min: 6.2 - max: 104.7). Fecal cortisol had no significant differences between males and females, but was higher in hunted (18.62 ng/g wet feces) than captured caribou (median 11.76 ng/g wet feces) in a subset of data with adult caribou from early spring 2018 (n=80, Z=-3.74, p<0.01). Among adult caribou captured in early spring, fecal cortisol varied significantly between years, H (df= 3, n= 135) = 49.6, p<0.01. Wilcoxon pairwise comparisons with a Bonferroni correction indicated that these yearly differences occurred between all years except for 2018 and 2021. Median fecal cortisol decreased over the years (2015: 41.2 ng/g of wet feces; 2016: 21.0 ng/g; 2018: 11.8 ng/g; 2021:13.34) (Figure 5).

### Results on health determinants

*We* detected exposure of DU caribou to all the pathogens tested (Table 2). Sample prevalence of antibody detection was different depending on the sampling method only for *Brucella*, which was higher in hunted caribou (34.3%; 95% CI: 20.8-50.8) than captured caribou (9.8%; 95% CI: 4.3-21.0) (n=86, p=0.01, Fisher’s exact test), assessed in a subset of data from adult caribou sampled in early spring 2018. Sample prevalence for pathogen exposure varied significantly by year using Fisher’s exact tests, for all pathogens except *Brucella* and Pestivirus (Table 2).

**Table 2.** Sample prevalence and confidence intervals for parasites and pathogens in adult Dolphin and Union caribou. These results include samples collected both by hunters and during caribou captures.

|  | 2015¶ | 2016¶ | 2017 | 2018 | 2019 | 2020 | 2021 | Total |
| --- | --- | --- | --- | --- | --- | --- | --- | --- |
| **Antibody detection – sample prevalence % (CI 95%)** |  |  |  |  |  |  |  |  |
| Pestivirus | 29.2<br>(14.9 – 49.2) | 21.7<br>(9.7 – 41.9) | 0.0<br>(0 – 39.0) | 23.6<br>(16.0 – 33.4) | 13.9<br>(6.6 – 27.3) | 15.4<br>(4.3 – 42.2) | 8.3<br>(2.9 – 21.8) | 18.8<br>(14.3 – 24.3) |
| α-herpesvirus* | 85.7<br>(68.5 – 94.3) | 100<br>(81.6 – 100) | NA | 83.1<br>(74.0 – 89.5) | 83.7<br>(70.0 – 91.9) | 15.4<br>(4.3 – 42.2) | 75.0<br>(58.9 – 86.2) | 79.6<br>(73.9 – 84.3) |
| *Brucella* sp.β | 10.7<br>(3.7 – 27.2) | 13.0<br>(4.5 – 32.1) | 0<br>(0 – 39.0) | 19.1<br>(12.3 – 28.5) | 20.9<br>(11.4 – 35.2) | 7.7<br>(0.4 – 33.3) | 13.5<br>(5.9-28.0) | 16.3<br>(12.2 – 21.5) |
| *Erysipelothrix rhusiopathiae* * | 21.4<br>(10.2 – 39.5) | 36.36<br>(19.7 – 57.0) | 33.3<br>(9.7 – 70.0) | 19.1<br>(12.3 – 28.5) | 6.9<br>(2.4 – 18.6) | 0<br>(0 – 22.8) | 16.2<br>(7.6 – 31.1) | 17.6<br>(13.3 – 23.0) |
| *Neospora caninum* * | 8.3<br>(2.3 – 25.8) | 33.3<br>(16.3 – 56.2) | NA | 2.2<br>(0.11 – 11.6) | 0.0<br>(0 – 48.9) | NA | 0.0<br>(0.0 – 9.4) | 7.0<br>(3.7 – 12.8) |
| **Parasite detection – sample prevalence % (CI 95%)** |  |  |  |  |  |  |  |  |
| *Besnoitia tarandi* (skin) | NA | 33.3<br>(1.7 – 79.2) | 40<br>(11.8 – 76.9) | 39.4<br>(24.7 – 56.3) | 43.2<br>(28.7 – 59.1) | NA | NA | 41.0<br>(30.8 – 52.1) |
| Prostrongylid Larvae/gram (feces) # | 5.9<br>(0.3 – 27.0) | 22.2<br>(9.0 – 45.2) | NA | 7.5<br>(3.5 – 15.4) | 11.54<br>(4.0 – 29.0) | NA | 9.1<br>(3.1 – 23.6) | 9.8<br>(6.2 – 15.1) |

¶ Part of the antibody detection results were previously published in (Carlsson et al. 2019) and (Aguilar et al.).
\* Differences between years were statistically significant.

We found significant differences of sample prevalence between adult males and females only for Herpesvirus exposure (males: 93.0% and females: 76.5%, n=223, Fisher’s exact test=0.018). Because the proportion of males and females sampled per year varied, we further confirm the sex differences in herpesvirus exposure with a subset of data that only included hunted caribou and excluded data from 2020 (males: 97.3%, females: 71.9%, n=94, Fisher’s exact test p<0.01), which was the only year in which the sample prevalence on herpesvirus was statistically different from other years (table 2).

*Besnoitia* cysts were detected in the metatarsus skin in 40.0% (95% CI: 31.1–49.6) of the caribou analyzed (n=105), with a median cyst density of 0.24 cysts/mm^2^ (min.:0.039 - max.:1.059). *Besnoitia* cyst detection in adult caribou did not vary significantly by sex or year (Table 2). Low intensities of protostrongylid larvae per gram of feces (min.: 0.20, median: 3.75, max.: 148.32) of the parasites *P. andersoni* and/or *V. eleguneniensis* were found in 9.9% (95% CI: 6.0 – 16.0) of the fecal samples analyzed (n=141). Protostrongylid fecal larvae detection in adult caribou in spring did not significantly vary by the method of sampling, sex or year (⋂=174, Table 2), and parasite counts were aggregated following a negative binomial distribution.

In the hair elements analysis, several elements had more than 40% (Na, Mg and K) or 90% (Cr and Co) of their values below the limit of quantification (LOQ) of the technique and no further assessments were possible. Among the heavy metals that are not trace minerals, As was not detected in any sample, Cd was detected in two samples above the LOQ (min: 0.26 ppm; max: 0.33 ppm) and Pb levels were above the LOQ in 57% of the samples (min: 0.02; median: 0.05; max: 50.8). There were no significant differences between sex of the animals in any of the elements analyzed, but there were significant differences in Fe between captured (median: 8.8) and hunted (median:10.6) caribou, in a subset of data that included adult caribou sampled in spring 2018 (n=85, Z=-2.78, p<0.01). Apart from selenium, there were significant differences in all hair trace minerals among years in the subset of data of adult caribou sampled in spring (n=129, Table 3).

**Table 3.**
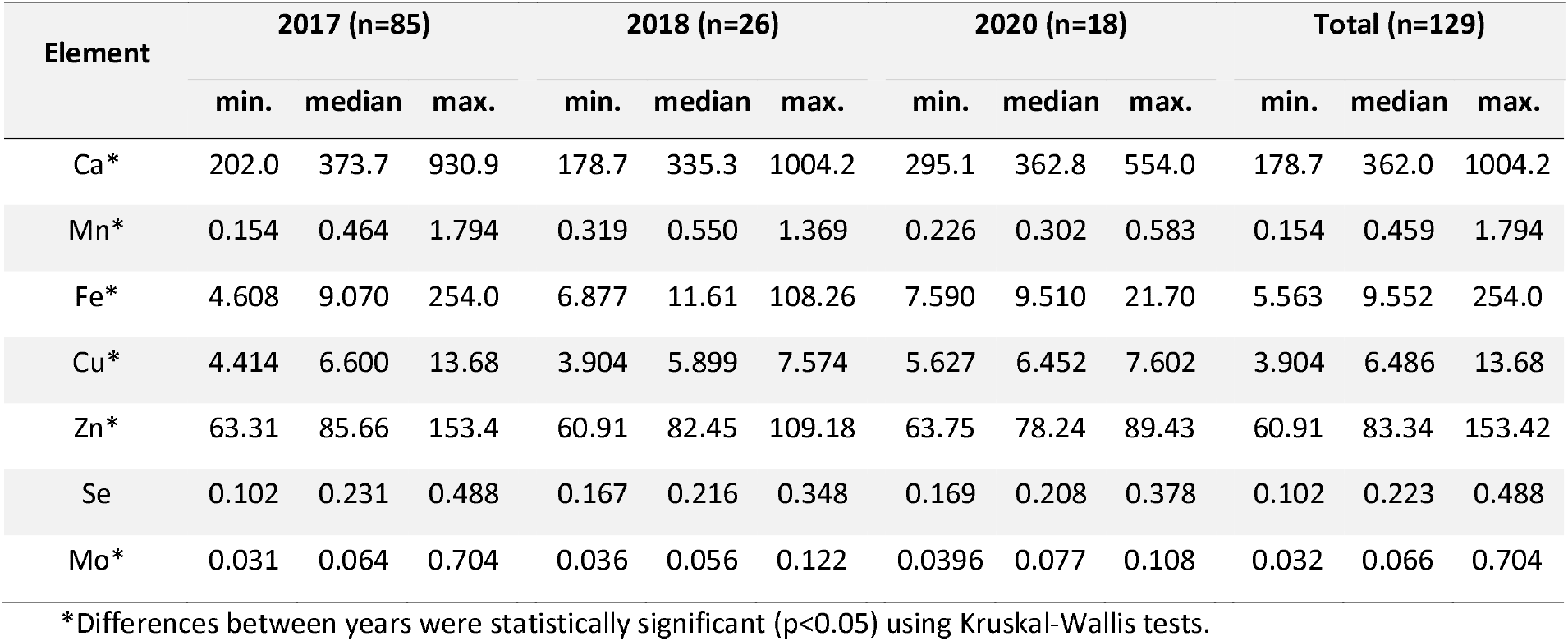
Concentration of trace minerals in parts per million (ppm) measured in hair from the neck of adult Dolphin and Union Caribou sampled in early spring. Element concentrations in hair are shown in the corresponding hair-growing year and not in the year in which the samples were collected.

### Abnormalities diagnosed during the sampling program

Between 2015 to 2021, eight cases of abnormal findings were submitted by hunters for pathological examination and were mostly diagnosed as brucellosis (Table 4). Among captured caribou, white-to-translucent spherical nodules in the eyes, consistent with *B. tarandi* cysts in the ocular sclera, were recorded in 8 out of 107 animals (not examined in 2015), and a one hindlimb hoof was overgrown in a cow captured in 2021 (negative to *Brucella* antibodies).

**Table 4.**
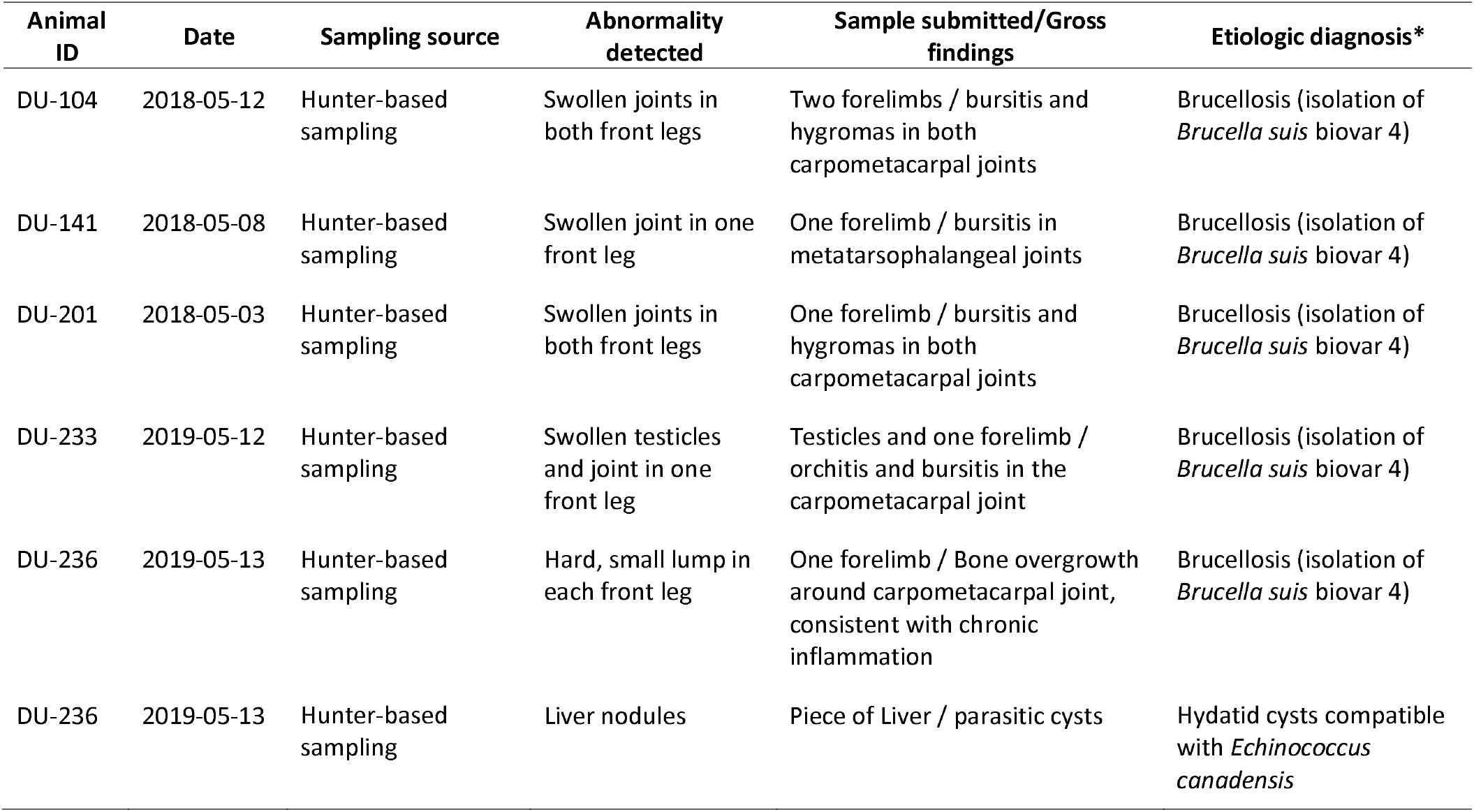
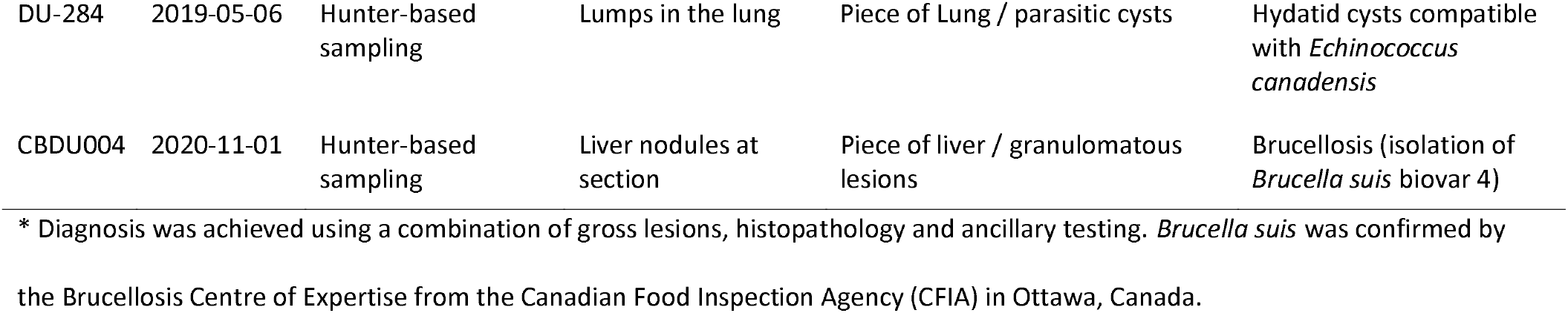
Cases of abnormalities in Dolphin and Union caribou submitted through the hunter-based sampling program between 2015 to 2021. These cases were submitted to the Diagnostic Service Unit, Faculty of Veterinary Medicine at the University of Calgary for diagnostic investigations by veterinary pathologists.

| Animal ID | Date | Sampling source | Abnormality detected | Sample submitted/Gross findings | Etiologic diagnosis* |
| --- | --- | --- | --- | --- | --- |
| DU-104 | 2018-05-12 | Hunter-based sampling | Swollen joints in both front legs | Two forelimbs / bursitis and hygromas in both carpometacarpal joints | Brucellosis (isolation of *Brucella suis* biovar 4) |
| DU-141 | 2018-05-08 | Hunter-based sampling | Swollen joint in one front leg | One forelimb / bursitis in metatarsophalangeal joints | Brucellosis (isolation of *Brucella suis* biovar 4) |
| DU-201 | 2018-05-03 | Hunter-based sampling | Swollen joints in both front legs | One forelimb / bursitis and hygromas in both carpometacarpal joints | Brucellosis (isolation of *Brucella suis* biovar 4) |
| DU-233 | 2019-05-12 | Hunter-based sampling | Swollen testicles and joint in one front leg | Testicles and one forelimb / orchitis and bursitis in the carpometacarpal joint | Brucellosis (isolation of *Brucella suis* biovar 4) |
| DU-236 | 2019-05-13 | Hunter-based sampling | Hard, small lump in each front leg | One forelimb / Bone overgrowth around carpometacarpal joint, consistent with chronic inflammation | Brucellosis (isolation of *Brucella suis* biovar 4) |
| DU-236 | 2019-05-13 | Hunter-based sampling | Liver nodules | Piece of Liver / parasitic cysts | Hydatid cysts compatible with *Echinococcus canadensis* |
| DU-284 | 2019-05-06 | Hunter-based sampling | Lumps in the lung | Piece of Lung / parasitic cysts | Hydatid cysts compatible with *Echinococcus canadensis* |
| CBDU004 | 2020-11-01 | Hunter-based sampling | Liver nodules at section | Piece of liver / granulomatous lesions | Brucellosis (isolation of *Brucella suis* biovar 4) |

\* Diagnosis was achieved using a combination of gross lesions, histopathology and ancillary testing. *Brucella suis* was confirmed by the Brucellosis Centre of Expertise from the Canadian Food Inspection Agency (CFIA) in Ottawa, Canada.

## Discussion

Wildlife health in its broad and modern definition, involves the capacity of animals to cope with stressors and maintain basic physiological functions, such as reproduction and survival (Ryser-Degiorgis 2013; Stephen 2014). Assessing wildlife health in remote regions requires a multi-faceted approach, including the use of diverse convenience sampling sources, and identifying and targeting relevant and informative indicators. We investigated multiple health indices in the endangered Dolphin and Union caribou herd over a 7-year period using samples from a collaborative health surveillance program that includes data and samples obtained by caribou harvesters and from capture events for monitoring purposes.

We classified our metrics on health into categories of outcomes, processes (host responses) and determinants (Pruvot et al.). This provides a practical framework to assess health in caribou when using multiple indicators, and complements previous frameworks that have been proposed (Wittrock et al. 2019; Peacock et al. 2020). Health outcomes such as reproduction, survival and body condition, are important indicators of caribou fitness which ultimately determine vital rates and population trends. As such, health outcomes are among the most important and immediate information to understand the trajectory of wild populations and relevant for guiding management decisions. However, to understand mechanisms and the fundamental drivers of the DU caribou decline, and to anticipate changes in population trajectory, it is important to examine health determinants and associated host responses in relation to these outcomes. This framework is broadly applicable, but is particularly valuable in remote wildlife populations from which population estimates and access to detailed examinations and investigations are limited.

### Health outcomes and populations trends

Our results suggest that body condition and pregnancy rates were unlikely the cause of the accelerated declines, rather, the decline was more likely linked to health determinants that affected adult survival, and perhaps calf survival. The overall pregnancy rate in DU caribou between 2015 and 2021, which includes the time of the steepest documented decline in this herd (between 2015 and 2018), was similar to other caribou herds from North America that were increasing or relatively stable (Bergerud 1980). Pregnancy rates and body condition from harvested DU caribou in these recent years, were similar to those documented in the late 80s, when the herd was increasing, and better than those in the early 2000s, when it had begun to decline (Figure 3, (Hanke et al. 2022)). In contrast, data from collared adult cows indicated a lower survival rate in the most recent years, 0.70 in 2015 and 0.62 in 2018 (Leclerc and Boulanger 2018; Leclerc and Boulanger 2020), when compared to the rate 0.76 obtained for the period 1999-2004 (Poole et al. 2010). Hunting by communities contributed to the DU caribou mortality, but when this cause was removed in survival analysis, the survival rate in 2015 remained low (0.72) and still consistent with a declining population (Leclerc and Boulanger 2018). Local knowledge of this herd indicates that the herd started to decline steadily around 2000s when skinnier animals and higher presence of diseased animals started to be noted (Tomaselli et al. 2018; Hanke et al. 2022). This is consistent with the pregnancy results recorded in early 2000s. However, the better pregnancy rates and body condition documented in the recent years suggest that different processes may have driven the steep decline between 2015 and 2018. Demographic studies on barren-ground herds found that herd size was most sensitive to adult survival (Haskell and Ballard 2007; Boulanger et al. 2011), and similar to our results, adult female survival was considered among the most important vital rate to explain the rapid decline of the Bathurst herd between 2006 and 2009 (Boulanger et al. 2011).

### Cortisol as a general host response to stressors

Cortisol concentrations in DU caribou hair and feces had a peak at the beginning of the study period and decreased over the years. Hair cortisol most likely reflected cumulative stressors during the main hair growth period (summer), or perhaps throughout the year (Rakic et al.), yet a significant decrease was documented. This pattern followed by both faecal and hair cortisol is coincident with the most accelerated declines of the DU caribou herd and the subsequent stabilization in the last population estimates between 2018 and 2020 at low numbers. Cortisol is a hormone that reflects the hypothalamic–pituitary–adrenal axis activity (Carlsson et al. 2016), and is a non-specific host response to stressors. It has been associated with different environmental stressors and disturbances in ungulates, including harvest and management practices, population density or demographics (Ewacha et al. 2017), but not with internal macroparasites (Carlsson et al. 2016). In DU caribou, higher allostatic loads seem to have been concurrent with the accelerated decline of the herd. Although this trend may be linked to the stressors that caused the fast herd decline, it could also be reflective of density-dependent mechanisms.

### Health determinants in relation to their effects and outcomes

*We* detected exposure to parasites (micro-and macro-) that can potentially affect caribou health either through: direct reproductive loss (*Brucella*, pestivirus, *N. caninum*, possibly α-herpesvirus); indirect effects through energetic costs (macroparasites), and; causing severe disease or death (mostly microparasites). Our results from health outcomes suggest that determinants that affect survival may have been more important in the most recent period. Therefore, we focus our assessments on microparasites that are more likely to cause severe health effects or death in caribou. However, it should be noted that antibody detection of pathogens that cause significant mortality after exposure may lead to an underestimation of its circulation, as positive animals would only be detected if they had recovered or survived from the infection.

The DU caribou had higher antibody prevalence against of *Brucella* and *α-herpesvirus* compared to recent estimates in other tundra caribou herds (Ducrocq et al. 2013; Carlsson et al. 2019). The effects of these pathogens in *Rangifer* are described in Table 1. *Brucella suis* biovar 4 was the main infectious agent isolated from abnormalities submitted by hunters during the study period (Table 4). Although sporadic, these lesions are consistently reported in caribou herds affected by brucellosis (Aguilar et al. 2022), and we can reasonably assume that at least a fraction of the exposed caribou develop severe lesions that could negatively impact survival. Harvesters in the 1990’s and early 2000s began reporting clinical signs consistent with brucellosis in the DU caribou herd (Tomaselli et al. 2018; Hanke et al. 2022). This corresponded with the initial decline of the herd and the lower pregnancy rates recorded in 2001-2003 (this work). While brucellosis may have contributed to the herd decline in earlier years, the overall good pregnancy rates and body condition suggest that additional health determinants may have been involved during the period of accelerated decline of the herd between 2015 and 2018.

*Erysipelothri× rhusiopathiae* has recently been identified as the cause of several mortality events in muskoxen (*Ovibos moschatus*), and has also been isolated from caribou carcasses (Forde et al. 2016a). Exposure in tundra caribou is common, and summer seroprevalence is positively associated with a range of environmental factors (Aleuy et al. 2022). The seroprevalence in DU caribou was higher in 2016 when the herd suffered its biggest decline and decreased in the subsequent years (Table 2, Figure 3). Increasing evidence indicates that a highly virulent strain of *E. rhusiopathiae* is circulating in the Canadian Arctic (Kutz et al. 2015; Forde et al. 2016a), however, given the opportunistic nature of this pathogen (Forde et al. 2016a), it cannot be discarded that its transmission and impact may also be driven by other stressors. While we cannot infer conclusive relationships of *E. rhusiopathiae* and health outcomes on the DU caribou herd, this pathogen is a significant cause of mortality and population declines in muskoxen (Kutz et al. 2015), is shared across multiple species (Forde et al. 2016b), and could also be involved in caribou mortalities.

The remaining pathogens studied by serology, including pestivirus, *N. caninum* and α-herpesvirus, are less likely to cause significant mortality and are mostly associated with reproductive loss or other type of syndromes (Table 1). However, sporadic mortalities related to increased virulence in circulating pestivirus strains have been documented in wild ungulates (Serrano et al. 2015; Colom-Cadena et al. 2019). It is noteworthy the decreasing trend in sample prevalence of pestivirus, *E. rhusipathiae, N. caninum* and α-herpesvirus along with the herd decline (Table 2). These trends may be caused by the reduction of the herd size and density-dependent mechanisms, decreasing their transmission or exposure due to smaller caribou aggregations. However, the peak and subsequent decrease of α-herpesvirus seroprevalence, could be triggered by other stressors that reactivated viral excretion and transmission during the accelerated decline (das Neves et al. 2009), which is supported by cortisol trends (Figure 5). The prevalence and intensity of *Besnoitia tarandi* cysts and protostrongylid larvae were at the low range of that reported in barren-ground caribou herds (Ducrocq et al. 2012).

There are no reference values for hair trace minerals in caribou and we can only compare our results with those from other *Rangifer* populations. Most element concentrations in the DU caribou were lower than those in a domestic reindeer herd from the University of Calgary (unpublished results, neck location n=13, medians Ca: 1565.51; Mn:2.62; Fe:34.53; Zn:114.25; Se:0.52; Mo: 0.24 ppm), and Se was lower than the Bluenose East herd (comparison only with neck location, Se:0.39 ppm) (Rakic et al.). In a northern contaminants study of the DU caribou, Cu and Se in kidneys of bulls were lower than those in other barren-ground caribou herds (Gamberg 2017). Together these data suggest that some elements may be lower in the range of DU caribou. Although micronutrient deficiencies are unlikely to directly cause mortality events, they may affect the resilience against other stressors.

### Methods, biases and sources of variation

Convenience sampling entails a nonrandom selection criterion of the animals. These sampling schemes, as compare to stratified random sampling, may suffer from selection or detection biases because are less likely to incorporate the variability introduced from biology and spatial and temporal scales (Conner et al. 2000; Nusser et al. 2008). Selection biases towards healthier animals may happen in both live-captured and harvested animals. Caribou captures were performed to study movements and space use, thus diseased or weak animals were unlikely to be selected. Similarly, Inuit harvesters prefer caribou with good body condition for consumption (Lyver and Gunn 2004). We found differences in pregnancy rates, fecal cortisol, and exposure to *Brucella* between captured and harvested animals, suggesting that the selection criteria between these two sampling sources may differ. We can expect that selection criteria in caribou captures performed from a helicopter is less context-dependent than hunter-based sampling, in which the hunter may not have as much opportunity to choose. In fact, clinical brucellosis with skinnier animals and swollen joints were only detected in caribou sampled by hunters (Table 4). Previous studies also found that herd vital rates obtained from collared caribou were better than they should be to explain the herd size trends, as derived from demographic models (Boulanger et al. 2011).

The higher fecal cortisol in harvested caribou from early spring may be associated with the concentration of hunters on specific areas from the DU migratory route (Figure 2). These intensive (few weeks) and geographically concentrated hunting efforts, may lead to repeated stress in caribou groups crossing these areas. We also found higher concentration of iron in hair from hunted caribou, but this finding is unexpected and difficult to explain. Other sources of variation in our metrics included the year of sampling for most of the metrics and sex in herpesvirus exposure, which has previously been reported in *Rangifer* (Lillehaug et al. 2003). However, samples from other seasons than spring and from males were poorly represented in our dataset and require further assessments.

### Conclusions

Data on health outcomes suggest that recent declines of the Dolphin and Union caribou herd were not associated with reduced pregnancy rates or poor body condition in early spring, rather, lower survival may have played a more important role. These results were also consistent when putting them into a broader context of herd population dynamics, by drawing on health data from historical caribou harvests and Indigenous knowledge of the herd. We provide insights into possible determinants and mechanisms that may have affected survival during the steep population decline (i.e. *Brucella suis* biovar 4, *Erysipelothri× rhusiopathiae* and lower trace minerals), yet complimentary approaches and targeted studies are needed to improve detection and investigations on caribou mortalities, for example, from collared animals.

The synergistic approach of harvester-based sampling and captures provided an increased sample size and different types of complementary information over different spatial and temporal scales, which is critical given the high individual, seasonal and yearly variability of health parameters in tundra caribou. With either method alone, the interpretation and understanding of the herd’s health would be limited, yet convenience sampling methods entail potential biases that need to be considered when analyzing these data. The widespread tundra caribou declines urgently require us to achieve a better understanding of the various mechanisms and determinants causing these declines. Surveillance strategies that combine captures, community-based sampling, participatory approaches, and local and Indigenous knowledge can greatly contribute to advance our knowledge on caribou and wildlife health.

## Acknowledgements

We thank the communities of Kugluktuk, Cambridge Bay and Ulukhaktok, particularly those hunters and residents that supported, participated, and together made possible the monitoring and sampling program. Thank you to Amanda Dumond, Bessie Inuktalik, and Beverly Maksagak from the Hunter’s and Trapper’s Associations, wildlife officers Russell Akeeagok and Allen Niptanatiak, and Brandon Langan, Terry Milton and Allen Pogotak who helped coordinate the program in community. We thank Andrea Hanke, Anja Carlsson, Angela Schneider and James Wang in the Kutz Lab, Dayna Goldsmith, Samuel Sharp, Melencio Nicolas, Jim Carlsen, and Betty Pollock in the Diagnostic Services Unit, Faculty of Veterinary Medicine at the University of Calgary, and David Kinniburgh of the Alberta Centre for Toxicology. Finally, we would like to thank to Don Russell who provided CARMA archived data on the Dolphin and Union caribou herd.

## References

Adams, B., J. Adamczewski, D. Cooley, G. Kofinas, R. Langvatn, R. Otto, D. Russell, R. White, et al. 2008. Rangifer Health & Body Condition Monitoring MANUAL.

Aguilar, X. F., I. H. Nymo, K. Beckmen, S. Dresvyanikova, I. Egorova, and S. Kutz. 2022. Brucellosis in the Arctic and northern regions. In Arctic One Health: challenges for arctic animals and people, ed. M. Tryland, 227–267. Cham, Switzerland: Springer.

Aguilar, X. F., F. Mavrot, O. Surujballi, L.-M. Leclerc, T. Davison, Hunter and Trapper’s Associations, M. Tomaselli, and S. Kutz. Brucellosis emergence in the Arctic, Canada. Under review in Emerging Infectious Diseases.

Albon, S. D., R. J. Irvine, O. D. D. Halvorsen, and R. Langvatn. 2017. Contrasting effects of summer and winter warming on body mass explain population dynamics in a food-limited Arctic herbivore: 1374–1389.

Aleuy, O. A., M. Anjold, K. Orsel, F. Mavrot, C. A. Gagnon, K. Beckmen, S. D. Côté, C. Cuyler, et al. 2022. Association of environmental factors with seasonal intensity of *Erysipelothrix rhusiopathiae* seropositivity among Arctic caribou. Emerging Infectious Diseases 28:1650–1658.

Ashley, N. T., P. S. Barboza, B. J. Macbeth, D. M. Janz, M. R. L. Cattet, R. K. Booth, and S. K. Wasser. 2011. Glucocorticosteroid concentrations in feces and hair of captive caribou and reindeer following adrenocorticotropic hormone challenge. General and Comparative Endocrinology 172: 382–391.

Ayroud, M., F. A. Leighton, and S. V Tessaro. 1995. The morphology and pathology of *Besnoitia* sp. in reindeer (*Rangifer tarandus tarandus*). Journal of Wildlife Diseases 31: 319–326.

Bergerud, A. T. 1980. A review of the population dynamics of caribou and wild reindeer in North America. In Proceedings of the Second International Reindeer and Caribou Symposium. 1979, ed. E. Reimers, E. Gaare, and S. Skjenneberg, 556–581. Trondheim, Norway: Direktoratet for vilt og ferskvannsfisk.

Boulanger, J., A. Gunn, J. Adamczewski, and B. Croft. 2011. A data-driven demographic model to explore the decline of the Bathurst caribou herd. Journal of Wildlife Management 75: 883–896.

Campbell, M., J. Ringrose, J. Boulanger, A. Roberto-Charron, K. Methuen, C. Mutch, T. Davison, and C. Wray. 2021. *An aerial Abundance estimate of the Dolphin and Union Caribou* (Rangifer tarandus groenlandicus x pearyi) *herd, Kitikmeot region, Nunavut–Fall 2020*. GN Technical report series – No□: 01-2021.

Carlsson, A. M., G. Mastromonaco, E. Vandervalk, and S. Kutz. 2016. Parasites, stress and reindeer: Infection with abomasal nematodes is not associated with elevated glucocorticoid levels in hair or faeces. Conservation Physiology 4:1–15.

Carlsson, A. M., P. Curry, B. Elkin, D. Russell, A. Veitch, M. Branigan, M. Campbell, B. Croft, et al. 2019. Multi-pathogen serological survey of migratory caribou herds: A snapshot in time. Plos One 14: e0219838.

Colom-Cadena, A., I. Marco, X. Fernández Aguilar, R. Velarde, J. Espunyes, R. Rosell, S. Lavín, and O. Cabezón. 2019. Experimental infection with high- and low-virulence strains of border disease virus (BDV) in Pyrenean chamois (*Rupicapra p. pyrenaica*) sheds light on the epidemiological diversity of the disease. Transboundary and Emerging Diseases 66:1619–1630.

Conner, M. M., C. W. McCarty, and M. W. Miller. 2000. Detection of bias in harvest-based estimates of chronic wasting disease prevalence in mule deer. Journal of Wildlife Diseases 36: 691–699.

COSEWIC. 2017. COSEWIC assessment and status report on the caribou, Dolphin and Union population, Rangifer tarandus, in Canada. Ottawa.

Curry, P. S., B. T. Elkin, M. Campbell, K. Nielsen, W. Hutchins, C. Ribble, and S. J. Kutz. 2011. Filter-paper blood samples for ELISA detection of *Brucella* antibodies in caribou. Journal of Wildlife Diseases 47:12–20.

Ducrocq, J., G. Beauchamp, S. Kutz, M. Simard, B. Elkin, B. Croft, J. Taillon, S. D. Côté, et al. 2012. Comparison of gross visual and microscopic assessment of four anatomic sites to monitor *Besnoitia tarandi* in barren-ground caribou (*Rangifer tarandus*). Journal of Wildlife Diseases 48: 732–738.

Ducrocq, J., G. Beauchamp, S. Kutz, M. Simard, J. Taillon, S. D. Côté, V. Brodeur, and S. Lair. 2013. Variables associated with *Besnoitia tarandi* prevalence and cyst density in barrenground caribou (*Rangifer tarandus*) populations. Journal of Wildlife Diseases 49: 29–38.

Ewacha, M. V. A., J. D. Roth, W. G. Anderson, D. C. Brannen, and D. L. J. Dupont. 2017. Disturbance and chronic levels of cortisol in boreal woodland caribou. Journal of Wildlife Management 81:1266–1275.

Forde, T. L., K. Orsel, R. N. Zadoks, R. Biek, L. G. Adams, S. L. Checkley, T. Davison, J. De Buck, et al. 2016a. Bacterial genomics reveal the complex epidemiology of an emerging pathogen in arctic and boreal ungulates. Frontiers in Microbiology 7:1–14.

Forde, T. L., R. Biek, R. Zadoks, M. L. Workentine, J. De Buck, S. Kutz, T. Opriessnig, H. Trewby, et al. 2016b. Genomic analysis of the multi-host pathogen *Erysipelothrix rhusiopathiae* reveals extensive recombination as well as the existence of three generalist clades with wide geographic distribution. BMC Genomics 17. BMC Genomics: 1–15.

Gamberg, M. 2017. Arctic Caribou Contaminant Monitoring Program 2016-2017. Report. Whitehorse, Yukon.

Gombart, A. F., A. Pierre, and S. Maggini. 2020. A review of micronutrients and the immune system—Working in harmony to reduce the risk of infection. Nutrients 12: 236.

Gunn, A., and D. Russell. 2017. *Caribou & wild reindeer* (Rangifer tarandus): *a terrestrial Focal Ecosystem Component in the Arctic – update for CBMP writing workshop March 2017*. CARMA.

Gunn, A., T. Leighton, and G. Wobeser. 1991. Wildlife diseases and parasites in the Kitikmeot region, 1984-90. File Report N°104. Coppermine, NWT: Department of Renewable Resources, Government of the Northwest Territories.

Hanke, A. N., M. Angohiatok, L.-M. Leclerc, C. Adams, and S. Kutz. 2022. A caribou decline foreshadowed by Inuit in the Central Canadian Arctic: a retrospective analysis. Arctic 74: 437–455.

Haskell, S. P., and W. B. Ballard. 2007. Modeling the Western Arctic caribou herd during a positive growth phase: potential effects of wolves and radiocollars. Journal of Wildlife Management 71: 619–627.

Hughes, J., S. D. Albon, R. J. Irvine, and S. Woodin. 2009. Is there a cost of parasites to caribou? Parasitology 136: 253–265.

Hunter, C. M., H. Caswell, M. C. Runge, E. V. Regehr, S. C. Amstrup, and I. Stirling. 2010. Climate change threatens polar bear populations: A stochastic demographic analysis. Ecology 91: 2883–2897.

Kafle, P., L. M. Leclerc, M. Anderson, T. Davison, M. Lejeune, and S. Kutz. 2017. Morphological keys to advance the understanding of protostrongylid biodiversity in caribou (*Rangifer* spp.) at high latitudes. International Journal for Parasitology: Parasites and Wildlife 6: 331–339.

Kafle, P., P. Peller, A. Massolo, E. Hoberg, L. M. Leclerc, M. Tomaselli, and S. Kutz. 2020. Range expansion of muskox lungworms track rapid arctic warming: implications for geographic colonization under climate forcing. Scientific Reports 10. Nature Publishing Group UK: 1–15.

Kutz, S., and M. Tomaselli. 2019. “Two-eyed seeing” supports wildlife health. Science 364: 1135–1137.

Kutz, S., J. Ducrocq, C. Cuyler, B. Elkin, A. Gunn, L. Kolpashikov, D. Russell, and R. G. White. 2013. Standardized monitoring of *Rangifer* health during International Polar Year. Rangifer 33:91.

Kutz, S., T. Bollinger, M. Branigan, S. Checkley, T. Davison, M. Dumond, B. Elkin, T. Forde, et al. 2015. *Erysipelothrix rhusiopathiae* associated with recent widespread muskox mortalities in the Canadian Arctic. Canadian Veterinary Journal 56: 560–563.

Kutz, S. J., E. P. Hoberg, L. Polley, and E. J. Jenkins. 2005. Global warming is changing the dynamics of Arctic host-parasite systems. Proceedings of the Royal Society B: Biological Sciences 272: 2571–2576.

Kutz, S. J., S. Laaksonen, K. Asbakk, and A. C. Nilssen. 2019. Parasitic infections and diseases. In Reindeer and Caribou: Health and Disease, ed. M. Tryland and S. J. Kutz, First, 177–235. Boca Raton, FL: CRC Press, Taylor and Francis.

Larska, M. 2015. Pestivirus infection in reindeer (*Rangifer tarandus*). Frontiers in Microbiology 6:1–5.

Leclerc, L.-M., and J. Boulanger. 2018. *Fall Population Estimate of the Dolphin and Union Caribou herd* (Rangifer tarandus groenlandicus × pearyi) *Victoria Island, October 2015 and Demographic population indicators 2015-2017*. Status report 2018-XX. Kugluktuk, NU.

Leclerc, L.-M., and J. Boulanger. 2020. *Population Estimate of the Dolphin and Union Caribou herd* (Rangifer tarandus groenlandicus × pearyi). *Coastal Survey, October 2018 and Demographic Indicators*. Kugluktuk, NU.

Lillehaug, A., T. Vlkøoren, I. L. Larsen, J. Åkerstedt, J. Tharaldsen, and K. Handeland. 2003. Antibodies to ruminant alpha-herpesviruses and pestiviruses in Norwegian cervids. Journal of Wildlife Diseases 39: 779–786.

Lyver, P. O. B., and A. Gunn. 2004. Calibration of hunters’ impressions with female caribou body condition indices to predict probability of pregnancy. Arctic 57: 233–241.

Macbeth, B., and S J Kutz. 2019. Rangifer Health a Holistic Perspective. In Reindeer and Caribou Health and Disease, ed. M. Tryland and Susan J Kutz, First Edit, 91–105. CRC Press.

Mallory, C. D., and M. S. Boyce. 2018. Observed and predicted effects of cliate change on Arctic caribou and reindeer. Environmental Reviews 26:13–25.

Morden, C.-J. C., R. B. Weladi, E. Ropstad, E. Dahl, O. Holand, G. Mastromonaco, and M. Nieminen. 2011. Fecal hormones as a non-invasive population monitoring method for reindeer. The Journal of Wildlife Management 75:1426–1435.

Myers-Smith, I. H., B. C. Forbes, M. Wilmking, M. Hallinger, T. Lantz, D. Blok, K. D. Tape, M. Maclas-Fauria, et al. 2011. Shrub expansion in tundra ecosystems: Dynamics, impacts and research priorities. Environmental Research Letters 6.

Neiland, K. A., J. A. King, B. E. Huntley, and R. O. Skoog. 1968. The diseases and parasites of Alaskan wildlife populations, Part I. Some observations on brucellosis in caribou. Bulletin of the Wildlife Disease Association 4: 27–36.

das Neves, C. G., T. Mørk, J. Thiry, J. Godfroid, E. Rimstad, E. Thiry, and M. Tryland. 2009. Cervid herpesvirus 2 experimentally reactivated in reindeer can produce generalized viremia and abortion. Virus Research 145: 321–328.

das Neves, C. G., S. Roth, E. Rimstad, E. Thiry, and M. Tryland. 2010. Cervid herpesvirus 2 infection in reindeer: A review. Veterinary Microbiology 143. Elsevier B.V.: 70–80.

Nusser, S. M., W. R. Clark, D. L. Otis, and L. Huang. 2008. Sampling considerations for disease surveillance in wildlife populations. Journal of Wildlife Management 72: 52–60.

Pachkowski, M., S. D. Côté, and M. Festa-Bianchet. 2013. Spring-loaded reproduction: Effects of body condition and population size on fertility in migratory caribou (*Rangifer tarandus*). Canadian Journal of Zoology 91: 473–479.

Parker, K. L., P. S. Barboza, and M. P. Gillingham. 2009. Nutrition integrates environmental responses of ungulates. Functional Ecology 23: 57–69.

Peacock, S. J., F. Mavrot, M. Tomaselli, A. Hanke, R. Nathoo, O. A. Aleuy, J. Di Francesco, F. Aguilar, et al. 2020. Linking co-monitoring to co-management: bringing together local, traditional, and sientigic knowldge in a wildlife status assessment framework. Arctic Science 6: 247–266.

Poole, K. G., A. Gunn, B. R. Patterson, and M. Dumond. 2010. Sea ice and migration of the Dolphin and Union caribou herd in the Canadian Arctic: an uncertain future. Arctic Institute of North America 63: 414–428.

Post, E., T. R. Christensen, R. A. Ims, E. Jeppesen, J. Madsen, A. D. Mcguire, and S. Rysgaard. 2009. Ecological dynamics across the arctic associated with recent climate change. Science 325: 1355–1358.

Post, E., B. A. Steinman, and M. E. Mann. 2018. Acceleration of phenological advance and warming with latitude over the past century. Scientific Reports 8. Springer US: 1–8.

Pruvot, M., X. Fernandez Aguilar, and S. Kutz. Wildlife Health Integrated Framework: A synthetic and operational framework to support transdisciplinary research and management. Under review.

R Development Core Team. 2017. R:A language and environment for statistical computing. Vienna: R Foundation for Statistical Computing. http://www.R-project.org.

Rakic, F. 2022. Hair biomarkers to support barren-ground caribou health monitoring and management. Master thesis. University of Calgary.

Rakic, F., X. Fernandez-Aguilar, M. Pruvot, D. P. Whiteside, G. Mastromonaco, L.-M. Leclerc, N. Jutha, and S. Kutz. Variation of hair cortisol in two herds of migratory caribou (*Rangifer tarandus):* implications for health monitoring. Conservation Physiology.

Russell, D. E., A. Gunn, and S. Kutz. 2018. Migratory tundra caribou and wild reindeer. In Arctic Report Card 2018, ed. E. Osborne, J. Richter-Menge, and M. Jeffries, 67–73.

Ryser-Degiorgis, M.-P. 2013. Wildlife health investigations: needs, challenges and recommendations. BMC veterinary research 9. BMC Veterinary Research: 223.

Schindelin, J., I. Arganda-Carreras, E. Frise, V. Kaynig, M. Longair, T. Pietzsch, S. Preibisch, C. Rueden, et al. 2012. Fiji: An open-source platform for biological-image analysis. Nature Methods 9: 676–682.

Serrano, E., A. Colom-Cadena, E. Gilot-Fromont, M. Garel, O. Cabezón, R. Velarde, L. Fernández-Sirera, X. Fernández-Aguilar, et al. 2015. Border disease virus: an exceptional driver of chamois populations among other threats. Frontiers in Microbiology 6:1307.

Stephen, C. 2014. Toward a modernized definition of wildlife health. Journal of Wildlife Diseases 50:427–430.

Tomaselli, M. 2022. Participatory epidemiology and surveillance for wildlife health. In Wildlife Population Health, ed. C. Stephen, 49–63. Springer, Cham.

Tomaselli, M., S. Kutz, C. Gerlach, and S. Checkley. 2018. Local knowledge to enhance wildlife population health surveillance: Conserving muskoxen and caribou in the Canadian Arctic. Biological Conservation 217. Elsevier: 337–348.

Wittrock, J., C. Duncan, and C. Stephen. 2019. A determinants of health conceptual model for fish and wildlife health. Journal of Wildlife Diseases 55: 285–297.

Zalatan, R., A. Gunn, and G. H. R. Henry. 2006. Long-term abundance patterns of barren-ground caribou using trampling scars on roots of Picea mariana in the Northwest Territories, Canada. Arctic, Antarctic, and Alpine Research 38: 624–630.

