## Supplementary Information for "An integrative and multi-indicator approach for wildlife health applied to an endangered caribou herd"

### Supplementary Information S1. Methods on the study area, the caribou population and sampling methods

#### *1.1 The Dolphin and Union caribou herd and its population trends*

The Dolphin and Union (DU) caribou is a migratory tundra caribou herd endemic to the Canadian Arctic, and is considered a separate Designatable Unit by the Committee on the Status of Endangered Wildlife in Canada (COSEWIC) due to its unique genetics, morphology, and ecology (COSEWIC 2017). This herd ranges in a vast area that covers part of the Kitikmeot and the Inuvialuit Arctic regions from Nunavut (NU) and Northwest Territories (NWT) in Canada. It migrates seasonally from its wintering grounds on the mainland to its summer range on Victoria Island, where it calves. Its southern range may overlap with barren-ground caribou (*R. tarandus groenlandicus*) and on its northern range with Peary caribou (*R. tarandus pearyi*) in some years (COSEWIC 2017). The herd may have once numbered 100,000 in the early 20th century (COSEWIC 2017), but has historically fluctuated and has been declining since the 1990s or 2000s according to local knowledge (Tomaselli et al. 2018; Hanke et al. 2022). Tracking trends in abundance is limited because there were only five aerial surveys from 1997 to 2020. The herd size in 1997 was estimated 34,558 animals (95% CI: 27,757-41,359) (Dumond and Lee 2013), and it has since declined by more than 88% to the most recent estimate of 3,815 in 2020

(95% CI: 2,930–4,966) (Campbell et al. 2021). The steepest decline was recorded between the estimates of 2015 (18,413; 95% CI: 11,644-25,182) and 2018 (4,105; 95% CI = 2,931-5,750), representing a reduction of 78% in herd size (Leclerc and Boulanger 2020). In 2017, COSEWIC re-assessed the DU herd from Special Concern to Endangered (COSEWIC 2017).

*1.2. Sampling methods and research permits*

In both community-based sampling and live-captured caribou, sampling methods were standardized following modified protocols of the CircumArctic Rangifer Monitoring and Assessment Network (CARMA) (Kutz et al. 2013). For the community-based sampling, kits with materials and instructions were provided to the hunters before their field trips. These kits consisted in a form to fill with basic information of the animal and the hunting event, and envelopes or bags for the independent collection of samples: blood in Nobuto filter paper strips (Toyo Roshi Kaisha, Ltd., Tokyo, Japan), feces, hair from the area of the neck, the left kidney with the surrounding fat, the lower jaw, the left hind leg and any abnormality observed by the hunter (Kutz et al. 2013; Tomaselli and Curry 2019). In captured caribou, only the non-invasive samples from the aforementioned protocol were collected (blood, hair and feces). The only difference between the two sample sources is that hair samples of captured caribou were plucked versus collected from the piece of skin taken from hunter-harvested animals (Kutz et al. 2013). All samples were maintained frozen upon collection at ambient outdoor temperatures or in freezers and then stored at -20 ºC in freezers until laboratory analyses. Abnormalities found by hunters were kept frozen and directly submitted to the Diagnostic Services Unit, Faculty of Veterinary Medicine, University of Calgary for further pathological investigation.

Sample collections were performed under the research permits WL2016-58 and WL2019-51 from the Government of Nunavut and WL5004469, WL500664 and WL500877 from the Government of Northwest Territories. The animal use protocols for research on biological sampling were approved (AC18-0093) by the Animal Care Committee from the University of Calgary, following the current Guidelines of the Canadian Council on Animal Care.

**Supplementary Information S2. Methods on body condition**

Body condition was measured following the CARMA protocol (Adams et al. 2008). Body condition metrics in harvested animals included the percentage of metatarsus marrow fat, Kidney Fat index (as described by Riney 1955 (Riney 1955)), back fat depth measured in cm as the thickest fat along an incision cut cranially from a 45 degree angle from the tail head (Kutz et al. 2013) and a hunter qualitative assessment (Kofinas et al. 2003). In live-captured animals, a body condition score that range from 1 to 12 was derived from the palpation of different body parts (Adams et al. 2008).

**Supplementary Information S3. Methods on Caribou Fecal and Hair Hormone analyses**

*3.1 Fecal hormone extracts*

To extract steroid hormones from feces, fecal samples were homogenized (for even distribution of metabolites) and weighed for 0.50 ± 0.02g weighed. Then 5 ml of 80% methanol was added for ratio of 0.1g/ml (Carlsson et al. 2016). Samples were vortexed briefly and hormones extracted overnight (16-18 hours) on a rotator plate (MBI Lab Equipment orbital shaker, 100 rpm) at room temperature. The samples were then centrifuged (10 min at 2400 x g) and the supernatant (fecal extract) decanted. Fecal extracts were stored at -20°C until hormone analysis.

*3.2 Hair hormone extracts*

The follicles were cut off before the hair samples were washed. For washing, the hair was immersed in distilled water and rubbed by hand for 2 minutes in 669 ml plastic containers. Then the hair samples were dried in a paper towel. A portion of the washed hair was vortexed with 15 ml of distilled water for 10 seconds in 20ml glass vials, then soaked for 5 minutes. Liquid was removed and a second 15 ml distilled water was added, vortexed for 10 seconds and removed immediately. Finally, 15 ml of 100% methanol was added, vortexed for 10 seconds and removed immediately. The process was repeated if any surface contamination (e.g., blood) was noticeable. Fully washed hair was dried in paper towel and stored in envelopes at room temperature.

Washed and dried hair was cut in 5mm pieces and weighed into 7ml glass scintillation vials. Hair samples weighting between 0.015-0.055 g were extracted with 100% methanol for a ratio of

0.01 g of hair/ml of methanol, on a rotator plate (MBI Lab Equipment orbital shaker, 100 rpm) at room temperature for 24 hours. The samples were then centrifuged (5 min at 2400 x g) to pipet the supernatant (hair extract) and place it into new glass 7ml vials. Hair extracts were stored at -20°C until hormone analysis.

*3.3 Hormone analysis by Enzyme Immunoassay*

For cortisol analysis in fecal samples, 40 µl of the fecal extracts were evaporated in a fume hood and the dried extract reconstituted in 160 µl EIA buffer for a 1:4 dilution. For cortisol analysis in hair samples, 1500 µl of the hair extracts were evaporated in a fume hood and the dried extract reconstituted in 150 µl EIA buffer for a 10-fold concentration. Cortisol was measured using an Enzyme Immunoassay based on the protocols from C. Munro, UC Davis, previously described in Kummrow et al. 2011 and Majchrzak et al. 2015. Briefly, microtiter plates were coated with 50 µl of the hormone-specific antibody R4866 diluted at 1:10,250 in coating buffer (50 mm bicarbonate buffer, pH 9.6). The plates were incubated overnight at 4°C and washed with 0.15 M NaCl and 0.05% Tween 20. For the assay, the wells were loaded with 50 µL of hormone standards or reconstituted extracts along with 50 µL of horseradish peroxidase diluted 1: 33,400 in EIA buffer. After two-hour incubation at room temperature, the plates were washed and 100 µL substrate solution (ABTS) was added. The absorbance was measured at 405nm using a spectrophotometer (Epoch 2 microplate reader, BioTek, Winooski, VT, USA). All samples and standards were run in duplicate.

For progesterone analysis, fecal extracts were diluted 1:20 to 1:500 in EIA buffer (0.1 mM sodium phosphate buffer, containing 9 g of NaCl and 1 g of bovine serum albumin per litre, pH 7). Progesterone was then measured using an EIA based on the protocols C. Munro, UC Davis, previously described in Kummrow et al. 2011 and Majchrzak et al. 2015. The procedure similar to the methods used in the cortisol assay, but for progesterone the plates were coated with the progesterone-specific antibody CL425 diluted at 1:8,800 in coating buffer, and the wells were loaded with 50 µL of diluted extracts along with 50 µL of horseradish peroxidase diluted 1:40,000.

All fecal progesterone, fecal cortisol and hair cortisol from samples collected in 2018-2021 was analyzed at the Endocrinology Lab at the Toronto Zoo, and the cortisol from hair samples collected in 2015-2017 was measured at the University of Saskatchewan (Faculty of Veterinary Medicine).

**Supplementary Information S4. Methods on *Besnoitia tarandi* cyst detection and maximal cyst density calculations**

The methods used for *Besnoitia tarandi* detection and maximal cyst density count were previously described (Ducrocq et al. 2012). Briefly, a piece of 2x2 cm of the metatarsal skin was fixed in 10% buffered formalin for at least 48 hours. A transvers section was processed by routine methods and 4 µm-thick sections of paraffin-embedded tissues were stained with hematoxylin and eosin at the Diagnostic Services Unit, Faculty of Veterinary Medicine, University of Calgary. Using light microscopy, an American College of Veterinary Pathologists (ACVP) board-certified veterinary pathologist (JLR) assessed slides for the presence or absence of *Besnoitia* cysts. For samples in which cysts were present, the pathologist selected the area of highest cyst density and using Olympus cellSense software, captured an image with the 1.25X microscope objective. We counted the number of tissue cysts observed in this image, using the ImageJ free software to calculate the total skin surface area (Schindelin et al. 2012). The area used for the cyst count was the dermis, extending from the epidermis of the skin to the base of the hair follicles and adnexal structures (Ducrocq et al. 2012). We used these values to calculate the estimated number of *B. tarandi* cysts per square millimetre.

**Supplementary Information S5. Methods on element concentrations in caribou hair**

In hair samples submitted as a piece of hide from the neck, about 200 mg of hair was shaved with a razor blade at about 2-3 mm from the skin. All hair samples were washed with a series of 75ml of 95% Ethanol followed by 75ml of Type I ultrapure water at least three times or until no surface contamination (e.g., blood) was noticeable. The washed hair was air-dried for more than a week or oven dried at 50°C for 24 hours in paper envelopes. About 70 mg of the washed hair was weighed in a Teflon vial and 2 mL of 70% nitric acid ($HNO_3$, TraceMetal grade, Fisher Scientific) was added. Digestion was performed using a high-pressure microwave digester

(ETHOS EZ Microwave Digestion System, Milestone, Sorisole, Italy), the temperature was gradually increased from room temperature to 180 °C over 25 minutes and then held at 180 °C for 15 minutes. Each digestion run also included the digestion of standard reference material NIST SRM 2976 (freeze-dried mussel tissue, National Institute of Standards and Technology) and DORM 3 (fish protein, National Research Council Canada), as well as an acid blank control (2 mL $HNO_3$). After the digestion was completed and samples reached room temperature, each sample was transferred to Falcon polypropylene test tube (VWR), diluted to 4 ml with Type I water, and stored at 5°C until analysis.

Samples were further diluted 10X with Type I water before analysis. Elemental analysis was carried out using inductively coupled plasma mass spectrometry (Agilent 8800 Triple Quadrupole ICP-MS) at the Alberta Centre for Toxicology. Each analysis run included the analysis of controls NIST SRM 1640a (trace elements in natural water) and Multi-element Standard (SCP Science). The acceptable criteria for controls were ± 20% of the certified values for all elements. We quantified a panel of trace minerals and contaminants that included: calcium (Ca), chromium (Cr), magnesium (Mg), manganese (Mn), iron (Fe), cobalt (Co), copper (Cu), zinc (Zn), potassium (K), selenium (Se), sodium (Na) and molybdenum (Mo) and three contaminant heavy metals, arsenic (As), cadmium (Cd) and lead (Pb).

**Supplementary Information S6: Analyses for ancillary data**

Caribou sampled were confirmed to belong to the Dolphin and Union herd by the Wildlife Genetics International (Nelson, British Columbia) using 18 microsatellite markers that enable the discrimination of the genetic ancestry (Serrouya et al. 2012). Animals were assigned as having DU genetics when the microsatellite results from samples (feces or dried skin) had higher than 50% of DU profile (data not shown). The herd identity for each individual was decided based on information from their genetic ancestry and calving location (in capture-collared animals), or, in the absence of these data, on sampling location and morphological traits described by harvesters (Leclerc and Boulanger 2018).

The age of caribou was assessed by analyzing the cementum annuli from an incisor, when available (n=90) (Hamlin et al. 2000), or by the tooth eruption pattern of the incisors in the capture-released animals (Adams et al. 2008). An age class was assigned to all caribou; calf for <12 months, yearling for 13-24 months, subadult for 25-36 months and adult for 37 months or greater.

The age of caribou was assessed by analyzing the cementum annuli from an incisor, when available (n=90) (Hamlin et al. 2000), or by the tooth eruption pattern of the incisors in the capture-released animals (Adams et al. 2008). An age class was assigned to all caribou; calf for <12 months, yearling for 13-24 months, subadult for 25-36 months and adult for 37 months or greater.

**Table S1.** Dolphin and Union caribou sampled since 2015 in different monitoring programs, shown by year, location, and sampling source (hunted and captured animals). Animals that were sampled as Dolphin and Union but had barren-ground genetics are not included in total numbers but are shown separately in brackets.

| | 2015 | 2016 | 2017 | 2018 | 2019 | 2020 | 2021 | Total |
|---|---|---|---|---|---|---|---|---|
| Hunted Caribou – Indigenous communities | | | | | | | | |
| Cambridge Bay (NU) | 15* | 7* | 7* | 2* | 18 | 14 | NA | 63 |
| Kugluktuk (NU) | 0 | 0 | 0 | 40 | 49 (8) | 0 | 0 | 89 (8) |
| Ulukhaktok (NWT) | 0 | 0 | 0 | 2 | 2 | 1 | NA | 5 |
| Collared Caribou | | | | | | | | |
| Captures GN | 16 (9) | 18 (1) | 0 | 51 | 0 | 0 | 38 | 123 (10) |
| Total | 31 (9) | 25 (1) | 7 | 95 | 69 (8) | 15 | 38 | 280 (18) |

* Includes sport hunt; NU= Nunavut; NWT= Northwest Territories; GN=Government of Nunavut; NA=samples collected but not analyzed in this study.

**Table S2.** Tests used for detection of antibodies against selected pathogens in caribou samples and references of previous usage in *Rangifer*. All serological analyses were performed at the University of Calgary except for *Brucella* assays that were done at the National Brucellosis Reference Laboratory (Canadian Food Inspection Agency-Ontario Animal Health Laboratory, Ottawa).

| Target pathogen | Test (Manufacturer) | Type of test | References |
|---|---|---|---|
| Pestivirus | BVDV/MD/BDV p80 Ab Test (IDEXX Laboratories Inc., Maine, USA) | cELISA | (Carlsson et al. 2019) |
| Alpha-herspesvirus | SERELISA IBR/IPV gB Ab Mono Blocking (Synbiotics, Europe SAS, France)<br>Infectious Bovine Rhinotracheitis Virus (BHV-1) gB Antibody Test Kit (IDEXX Laboratories Inc., Maine, USA) | cELISA | (Das Neves et al. 2009; Carlsson et al. 2019) |
| *Brucella* | In-house competitive ELISA* | cELISA | (Gall et al. 2001; Curry et al. 2011) |
| *Erysipelothrix rhusiopathiae* | In-house indirect ELISA | iELISA | (Bondo et al. 2019; Aleuy et al. 2022) |
| *Neospora caninum* | *Neospora caninum* Antibody Test Kit, cELISA (VMRD Inc., Pullman, WA, USA) | cELISA | (Curry et al. 2014; Carlsson et al. 2019) |

cELISA: competitive Enzyme-linked immunoabsorbent assay; iELISA: indirect ELISA.
* Canadian Food Inspection Agency (CFIA), Brucellosis Centre of Expertise, Ottawa, ON, Canada.
¶ University of Calgary

### References


Adams, B., J. Adamczewski, D. Cooley, G. Kofinas, R. Langvatn, R. Otto, D. Russell, R. White, et al. 2008. *Rangifer Health & Body Condition Monitoring MANUAL*.

Aleuy, O. A., M. Anjold, K. Orsel, F. Mavrot, C. A. Gagnon, K. Beckmen, S. D. Côté, C. Cuyler, et al. 2022. Association of Environmental Factors with Seasonal Intensity of *Erysipelothrix rhusiopathiae* Seropositivity among Arctic Caribou. *Emerging Infectious Diseases* 28: 1650–1658.

Bondo, K. J., B. Macbeth, H. Schwantje, K. Orsel, D. Culling, B. Culling, M. Tryland, I. H. Nymo, et al. 2019. Health survey of Boreal caribou (*Rangifer tarandus caribou*) in northeastern

British Columbia, Canada. *Journal of Wildlife Diseases* 55: 2018-01–018. doi:10.7589/2018-01-018.

Campbell, M., J. Ringrose, J. Boulanger, A. Roberto-Charron, K. Methuen, C. Mutch, T. Davison, and C. Wray. 2021. *An Aerial Abundance Estimate of the Dolphin and Union Caribou (*Rangifer tarandus groenlandicus x pearyi*) Herd, Kitikmeot Region, Nunavut – Fall 2020. GN Technical Report Series – No : 01-2021*.

Carlsson, A. M., G. Mastromonaco, E. Vandervalk, and S. Kutz. 2016. Parasites, stress and reindeer: Infection with abomasal nematodes is not associated with elevated glucocorticoid levels in hair or faeces. *Conservation Physiology* 4: 1–15. doi:10.1093/conphys/cow058.

Carlsson, A. M., P. Curry, B. Elkin, D. Russell, A. Veitch, M. Branigan, M. Campbell, B. Croft, et al. 2019. Multi-pathogen serological survey of migratory caribou herds: A snapshot in time. *Plos One* 14: e0219838. doi:10.1371/journal.pone.0219838.

COSEWIC. 2017. *COSEWIC assessment and status report on the Caribou, Dolphin and Union population, Rangifer tarandus, in Canada.* Ottawa.

Curry, P. S., B. T. Elkin, M. Campbell, K. Nielsen, W. Hutchins, C. Ribble, and S. J. Kutz. 2011. Filter-paper blood samples for ELISA detection of *Brucella* antibodies in caribou. *Journal of Wildlife Diseases* 47: 12–20. doi:10.7589/0090-3558-47.1.12.

Curry, P. S., C. Ribble, W. C. Sears, W. Hutchins, K. Orsel, D. Godson, R. Lindsay, A. Dibernardo, et al. 2014. Blood collected on filter paper for wildlife serology: detecting antibodies to *Neospora caninum*, West Nile Virus and five bovine viruses in *Rangifer tarandus* subspecies. *Journal of Wildlife Diseases* 50: 297–307. doi:10.7589/2012-02-047.

Ducrocq, J., G. Beauchamp, S. Kutz, M. Simard, B. Elkin, B. Croft, J. Taillon, S. D. Côté, et al. 2012. Comparison of Gross Visual and Microscopic Assessment of Four Anatomic Sites To Monitor *Besnoitia tarandi* in Barren-Ground Caribou (*Rangifer tarandus*). *Journal of Wildlife Diseases* 48: 732–738. doi:10.7589/0090-3558-48.3.732.

Dumond, M., and D. S. Lee. 2013. Dolphin and union caribou herd status and trend. *Arctic* 66: 329–337. doi:10.14430/arctic4311.

Gall, D., K. Nielsen, L. Forbes, W. Cook, D. Leclair, S. Balsevicius, L. Kelly, P. Smith, et al. 2001.

Evaluation of the fluorescence polarization asssay and comparison to other serological assays for detection of brucellosis in cervids. *Journal of Wildlife Diseases* 37: 110–118.

Hamlin, K. L., D. F. Pac, C. A. Sime, R. M. DeSimone, and G. L. Dusek. 2000. Evaluating the Accuracy of Ages Obtained by Two Methods for Montana Ungulates. *Journal of Wildlife Management* 64: 441–449.

Hanke, A. N., M. Angohiatok, L.-M. Leclerc, C. Adams, and S. Kutz. 2022. A caribou decline foreshadowed by Inuit in the Central Canadian Arctic: a retrospective analysis. *Arctic* 74: 437–455. doi:10.14430/arctic73826.

Kofinas, G., P. Lyver, D. Russell, R. White, A. Nelson, and N. Flanders. 2003. Towards a protocol for community monitoring of caribou body condition. *Rangifer*: 43–52. doi:10.7557/2.23.5.1678.

Kummrow, M. S., C. Gilman, P. Mackie, D. A. Smith, and G. F. Mastromonaco. 2011. Noninvasive analysis of fecal reproductive hormone metabolites in female veiled chameleons (*Chamaeleo calyptratus*) by enzyme immunoassay. *Zoo Biology* 30: 95–115. doi:10.1002/zoo.20318.

Kutz, S., J. Ducrocq, C. Cuyler, B. Elkin, A. Gunn, L. Kolpashikov, D. Russell, and R. G. White. 2013. Standardized monitoring of Rangifer health during International Polar Year. *Rangifer* 33: 91. doi:10.7557/2.33.2.2532.

Leclerc, L.-M., and J. Boulanger. 2018. *Fall Population Estimate of the Dolphin and Union Caribou herd (*Rangifer tarandus groenlandicus *x* pearyi*) Victoria Island, October 2015 and Demographic population indicators 2015-2017. Status report 2018-XX*. Kugluktuk, NU.

Leclerc, L.-M., and J. Boulanger. 2020. *Population Estimate of the Dolphin and Union Caribou herd (*Rangifer tarandus groenlandicus x pearyi*). Coastal Survey, October 2018 and Demographic Indicators.* Kugluktuk, NU.

Majchrzak, Y. N., G. F. Mastromonaco, W. Korver, and G. Burness. 2015. Use of salivary cortisol to evaluate the influence of rides in dromedary camels. *General and Comparative Endocrinology* 211. Elsevier Inc.: 123–130. doi:10.1016/j.ygcen.2014.11.007.

Das Neves, C. G., M. Roger, N. G. Yoccoz, E. Rimstad, and M. Tryland. 2009. Evaluation of three commercial bovine ELISA kits for detection of antibodies against Alphaherpesviruses in reindeer (*Rangifer tarandus tarandus*). *Acta Veterinaria Scandinavica* 51: 1–10. doi:10.1186/1751-0147-51-9.

Riney, T. 1955. Evaluating condition of free-ranging red deer (*Cervus elaphus*), with special reference to New Zealand. *New Zealand Journal of Science and Technology, Sect B* 36: 429–463.

Schindelin, J., I. Arganda-Carreras, E. Frise, V. Kaynig, M. Longair, T. Pietzsch, S. Preibisch, C. Rueden, et al. 2012. Fiji: An open-source platform for biological-image analysis. *Nature Methods* 9: 676–682. doi:10.1038/nmeth.2019.

Serrouya, R., D. Paetkau, B. N. McLellan, S. Boutin, M. Campbell, and D. A. Jenkins. 2012. Population size and major valleys explain microsatellite variation better than taxonomic units for caribou in western Canada. *Molecular Ecology* 21: 2588–2601. doi:10.1111/j.1365-294X.2012.05570.x.

Tomaselli, M., and P. Curry. 2019. Widllife health and disease surveillance. In *The Veterinary Laboratory & Field Manual*, ed. S. C. Cork and R. W. Halliwell, Third Edit, 420–432. 5m Publishing.

Tomaselli, M., S. Kutz, C. Gerlach, and S. Checkley. 2018. Local knowledge to enhance wildlife population health surveillance: Conserving muskoxen and caribou in the Canadian Arctic. *Biological Conservation* 217. Elsevier: 337–348. doi:10.1016/j.biocon.2017.11.010.
